# Caspase inhibition mitigates tau cleavage and neurotoxicity in iPSC-induced neurons with the V337M *MAPT* mutation

**DOI:** 10.1101/2021.01.08.425912

**Authors:** Panos Theofilas, Chao Wang, David Butler, Dulce O. Morales, Cathrine Petersen, Brian Chin, Teddy Yang, Shireen Khan, Raymond Ng, Rakez Kayed, Celeste M. Karch, Bruce L. Miller, Jason E. Gestwicki, Li Gan, Sally Temple, Michelle R. Arkin, Lea T. Grinberg

**Author notes:** Co-senior author (Michelle R. Arkin, Lea T. Grinberg). **Correspondence:** 1. Lea T. Grinberg, MD, PhD; John Douglas French Alzheimer’s Foundation Endowed Professor. Associate Professor of Neurology and Pathology; University of California, San Francisco; 675 Nelson Rising Lane, 211B, Box 1207; San Francisco, CA 94158, 2. Michelle R. Arkin, PhD; Professor of Pharmaceutical Chemistry; co-Director, Small Molecule Discovery Center; School of Pharmacy, University of California San Francisco; Box 2552; 1700 4th Street. San Francisco, CA. 94143.

## Abstract

Tau post-translational modifications (PTMs) are associated with progressive tau accumulation and neuronal loss in tauopathies, including forms of frontotemporal lobar degeneration (FTLD) and Alzheimer’s disease (AD). Proteolytic cleavage of tau by active caspases, including caspase-6, represents an underexplored tau PTM implicated in tau pathology. Caspase-cleaved tau is toxic and prone to self-aggregation in experimental models. To elucidate the presence and temporal course of caspase activation, tau cleavage, and neuronal death, we generated two neoepitope monoclonal antibodies (mAbs) against caspase-6 tau proteolytic sites and cortical neurons from induced pluripotent stem cells (iPSCs) with the frontotemporal dementia (FTD)-causing V337M *MAPT* mutation. FTLD V337M *MAPT* and AD postmortem brains showed positivity for both cleaved tau mAbs as well as active caspase-6. Relative to isogenic wild-type *MAPT* controls, V337M *MAPT* neurons showed a time-dependent increase in pathogenic tau in the form of tau oligomers, caspase-cleaved tau, and p-tau. Accumulation of toxic tau species in 3-month V337M *MAPT* neurons also increased vulnerability to stress, which was pharmacologically rescued by caspase inhibition. We propose a model in which time-dependent accumulation of caspase-cleaved tau in V337M *MAPT* neurons promotes neurotoxicity that is reversed by caspase-6 inhibition. Caspase-cleaved tau may be a biomarker of tauopathy, and caspases could be viable targets for therapeutic intervention against tau pathogenesis in FTLD and other tauopathies.

**Significance:** The mechanisms leading to tau pathology in frontotemporal dementia (FTD) and Alzheimer’s disease (AD) remain elusive. Experimental studies in AD demonstrate that tau cleavage by active caspase-6 contributes to tau pathology since cleaved tau may be toxic and prone to self-aggregation. Yet, the role of caspase-cleaved tau in promoting toxicity and cell death is unclear. Here, we generated two neoepitope monoclonal antibodies against caspase-6 tau and evaluated tau cleavage in postmortem human brains, iPSC-induced cortical neurons with the FTD-causing V337M *MAPT* mutation, and isogenic wild-type *MAPT* controls. Our results demonstrate a time-dependent accumulation of caspase-cleaved tau and increased neurotoxicity in the mutant iNs that is reversed by caspase-6 inhibition. Caspases could be viable therapeutic targets against tau pathology in tauopathies.

## Introduction

Tau post-translational modifications (PTMs) represent pathological hallmarks of neurodegenerative diseases collectively known as tauopathies, including frontotemporal lobar degeneration (FTLD) and Alzheimer’s disease (AD) (1, 2). Considering the absence of effective treatments against tauopathies, therapeutic interventions that delay or impede tau pathogenesis represent an urgent medical need. Although tau hyperphosphorylation (p-tau) is one of few PTMs extensively investigated in tauopathies for its role in tau aggregation and neurofibrillary (NFT) pathology (3, 4), the less explored proteolytic cleavage of tau by active caspases has recently sparked much research interest. In experimental models, caspase-cleaved tau promotes tau aggregation, tau spreading, and neurotoxicity in tauopathies (5–8).

Caspases (cysteine-aspartic proteases) are proteolytic enzymes with well-defined roles in cell death and inflammation. Apoptotic caspases are broadly classified as either initiator caspases that activate their effector counterparts or effector/ executioner caspases that process downstream substrates and cause the destruction of the cell (9–11). Accumulating evidence support a key pathogenic role of the effector caspase-6 in major diseases including AD, Huntington disease, and ischemic stroke (12–15). In AD, active caspase-6 is associated with three stages of cellular dysfunction. Caspase-6 activity is linked to neuronal inflammation (16, 17) and apoptosis by processing key proteins required for cell homeostasis. Active caspase-6 also cleaves tau at confirmed and putative sites, potentially contributing to tau aggregation (8, 18, 19). Moreover, active caspase-6 colocalizes with NFT-positive neurons in AD patients’ brains (15, 18, 20, 21), promotes axonal degeneration in several cellular and *in vivo* animal models of AD (22, 23), and its levels inversely correlate with cognitive performance in aged individuals (24).

We previously demonstrated a positive linear correlation between AD neuropathological stages and increasing levels of active caspase-6 and p-tau colocalization in the locus coeruleus and dorsal raphe nucleus, brain regions showing the earliest vulnerability to AD-tau pathology (15). Active caspase-6 promotes NFTs by cleaving tau after aspartic acid (Asp) at multiple sites, including Asp421 (D421) near the C-terminus (8, 18, 25). D421, which can be cleaved by multiple caspases, including caspase-3, has a stronger seeding propensity than full-length tau and compose the core of PHFs in humans (26) and experimental models of AD (5, 27–29). In addition to the well-characterized D421 site, other tau cleavage sites were considered to have a role in disease pathogenesis. Specifically, the N-terminal D13 (19) and C-terminal D402 sites (24, 28, 30) are thought to be cleaved primarily by caspase-6 and are found to mediate tau aggregation and toxicity *in vitro.* Yet, these sites are underexplored compared to D421, mainly due to a lack of available monoclonal antibodies (mAbs). Currently, D13 is detected by the loss of N-terminal tau epitopes recognized by the antibodies Tau-12 and 5A6 (19), and D402 is detected using a polyclonal antibody (28). Considering that early tau modifications are critical for our understanding of tau evolution and disease development, the generation of improved neoepitope antibodies against novel tau cleavage sites will shed light on pathological tau processes and further propose caspases as possible therapeutic targets in tauopathies.

To elucidate the presence and temporal course of caspase activation, tau cleavage, and neuronal death in tauopathies, we generated induced cortical neurons (iNs) from induced pluripotent stem cells (iPSCs) with the FTD-causing V337M *MAPT* mutation (tau^V337M^) (31–34) and WT isogenic controls (tau^WT^). We evaluated tau proteolysis using newly synthesized neoepitope mAbs against caspase-cleaved tau at D13 and D402 sites. iPSCs and their subsequent conversion to iNs by expression of neuronal transcription factors offer a clinically-relevant model of human tau pathogenesis in a disease-specific genetic background (33, 35–38). Our results demonstrate a time-dependent accumulation of caspase-cleaved tau and increased neurotoxicity in the tau^V337M^ iNs that is reversed by caspase-6 inhibition. This study offers insights into the potential of caspase-6 and other caspases as targets for therapeutic intervention against tau pathology in FTD and other tauopathies.

## Results

### Generation of caspase-6 cleaved tau neoepitope monoclonal antibodies

Besides TauC3 (D421), mAbs that recognize specific caspase-cleaved tau epitopes are crucially missing. Here, we developed two neoepitope mAbs against D13 and D402 tau cleavage sites (Fig. 1a) using hybridoma technology. Neoepitope antibodies selectively bound to cleaved tau (1-402 or 14-441) but not full-length tau (1-441) as detected by enzyme-linked immunosorbent assay (ELISA) and western blot analyses (Fig. S1; S2-a, b; See Supplemental Experimental Procedures). Antibody specificity was also confirmed by immunofluorescence (IF) and antigen competition assays where mAb.D402.1 and mAb.D13.1 primary antibodies were preincubated with the cognate peptides used as immunogens (Table S1), resulting in loss of antibody signal (Fig. S2-c). Together with TauC3, mAb.D402.1, and mAb.D13.1 neoepitope antibodies were used to identify caspase-mediated pathological changes in the tau^V337M^ and tau^WT^ iNs by western blot protein analyses and multiplex IF.

**Fig. 1.**
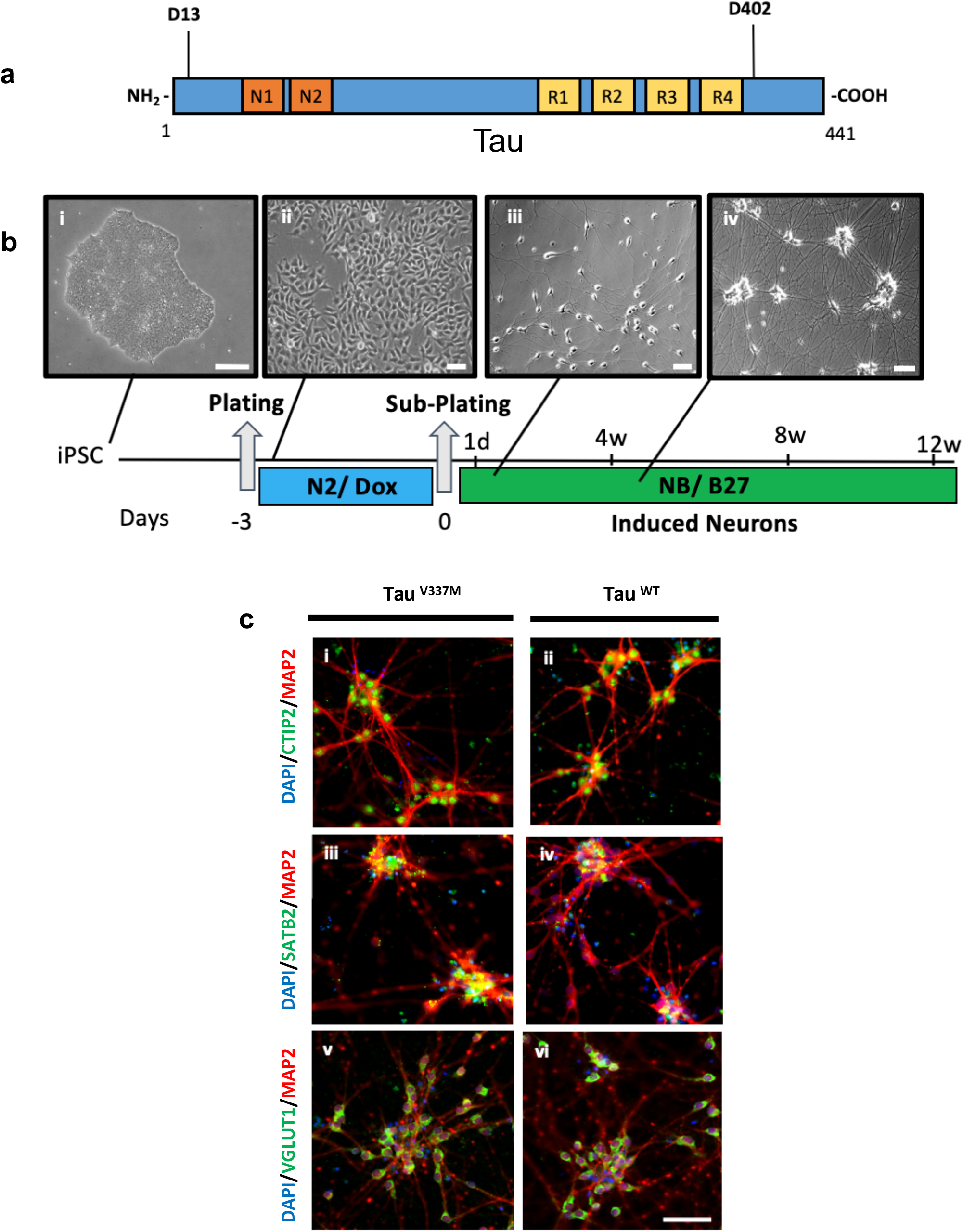
Generation of caspase-6 cleaved tau neoepitope antibodies and iPSC-induced neurons. (a) Schematic depicting two caspase-6 cleaved tau sites targeted by monoclonal neoepitope antibodies mAb.D13.1 (D13; 14-441) and mAb.D402.1 (D402; 1-402), at the N- and C-terminus of tau, respectively. (b) Timeline of iPSC differentiation to neurons following doxycycline-inducible expression of Ngn2. Phase-contrast images represent individual stages of the differentiation protocol, including iPSCs (i), neuronal precursors (ii), immature neurons (iii), and mature neurons (iv). (c) IF characterization of neurons at 4-weeks post differentiation. Mutant (tau^V337M^) and control (tau^m^) neurons were positive for cortical (green; i-iv), glutamatergic (green; v-vi), and neuronal (red; i-vi) markers. Nuclei were stained with DAPI (blue; i-vi). Scale bars: (b) i: 200 μm, ii-iv: 50 μm (c) 60 μm. See also Table S1 and Fig. S1-S3.

### Characterization of iPSC-induced neurons

Considering that V337M *MAPT* mutation (tau^V337M^) causes FTD in humans, we generated heterozygous tau^V337M^ iNs and isogenic tau^WT^ controls to examine the disease mechanisms involving caspase activation and cleaved tau pathology in a clinically relevant cell culture model of tauopathy. Neuronal induction of a well-characterized human tau^WT^ line was performed using TALEN-based integration of a doxycycline-inducible *Ngn2* transgene into the AAVS1 safe harbor as previously described (33, 39). CRISPR/Cas9 genome editing was used to introduce the tau^V337M^ into tau^WT^ iPSCs to study the mutation’s effects in isolation from the donor’s genetic background. Both iPSC groups had a normal karyotype (Fig. S3-a) and typical colony-type morphology (Fig. 1b). Moreover, genomic DNA sequencing of tau^V337M^ and tau^WT^ iPSCs confirmed the presence of the heterozygous tau^V337M^ in exon 12 (Fig. S3-b), and homogeneous expression of the pluripotency markers NANOG, OCT4, and SOX2 (Fig. S3-c) in the iNs confirmed the absence of NGN2 expression leakage without the addition of doxycycline. Following doxycycline treatment, iNs exhibited neuronlike morphology in 5-7 days and mature neuronal morphology between 3-4 weeks (Fig. 1b).

One month old iNs were positive for the microtubule-associated protein 2 (MAP2) neuronal marker in the cytoplasm and neurites, and the deep and upper cortical nuclear markers CTIP2 and SATB2, respectively, as confirmed by IF (Fig. 1c). iNs were also positive for the glutamatergic marker vesicular glutamate transporter 1 (VGlut1) in the cytoplasm and neurites, an anticipated outcome of the *NGN2* expression (33, 40). Overall, our findings demonstrate that at one month post-differentiation, iNs exhibit neuron-specific morphology and express cortical and glutamatergic markers.

### Increased levels of pathological tau in the tau^V337M^ neurons

Since FTLD-tau is characterized by progressive accumulation of toxic tau species and neuronal loss, we aimed to examine the presence and temporal course of tau pathological accumulation in tau^V337M^ in the mutant iNs relative to WT isogenic controls. We used a panel of tau antibodies, including total and oligomeric tau, 3R and 4R tau, and caspase-cleaved tau, to compare iNs cultured from 1 to 3 months by western blot (Fig. 2).

**Fig. 2.**
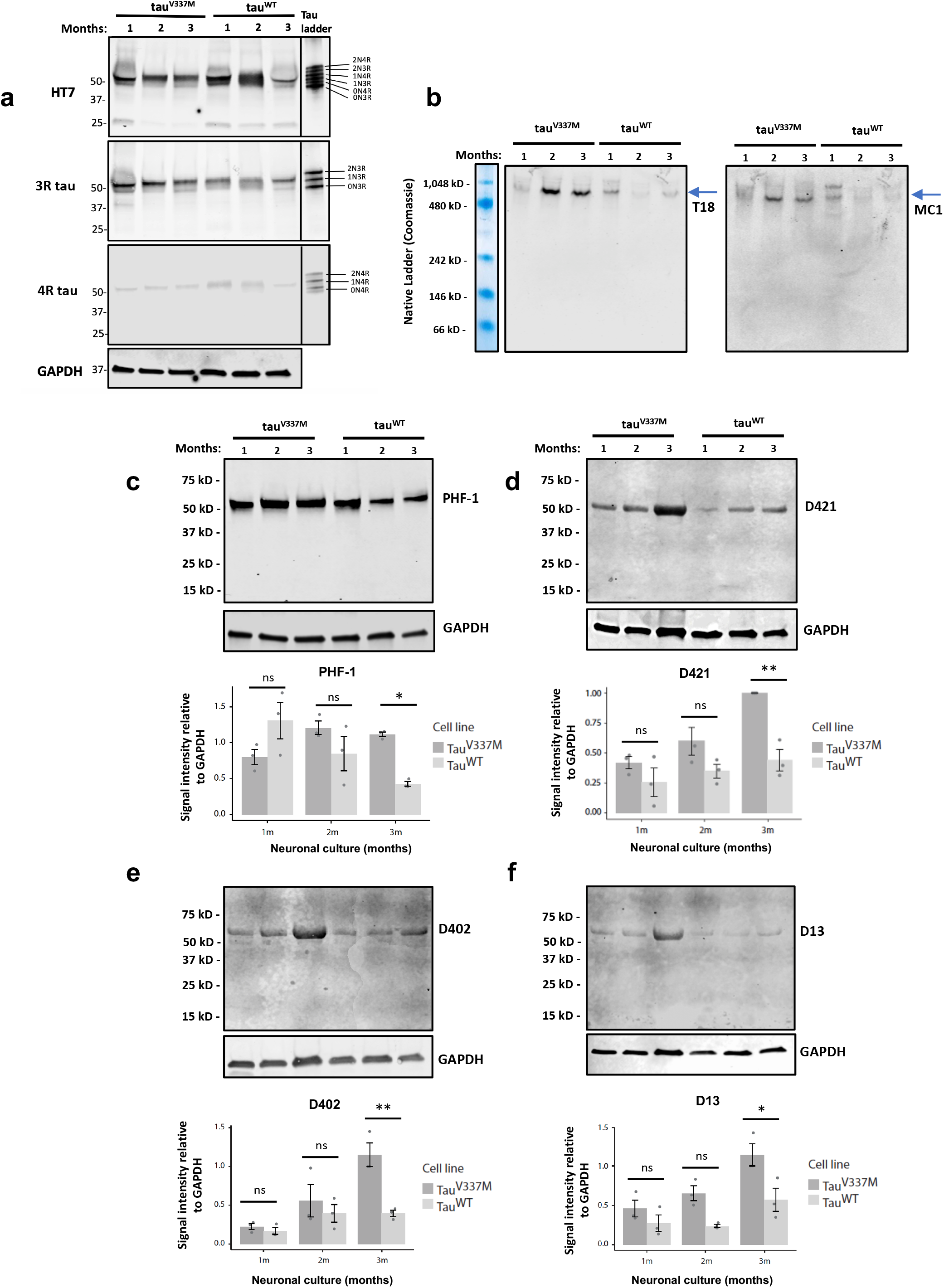
Time-dependent tau pathological changes in the tau^V337M^ neurons detected by western blot analyses. (a) Expression of total tau (HT7) and tau isoforms (3R and 4R tau) in tau^V337M^ and tau^m^ neurons cultured from 4 to 12 weeks. Antibody specificity was confirmed by a recombinant tau ladder containing all six tau isoforms (right panels). (b) Oligomeric (T18) and conformational (MC1) tau levels were detected under non-denaturing conditions for preserving protein conformation. The molecular weight of the proteins was estimated using a protein standard for native electrophoresis stained with Coomassie blue (far-left panel) for band visualization (see also Fig. S4). (c-f) Semi-quantification of p-tau (PHF-1) and caspase-cleave tau (D421, D402, and D13) protein levels based on band intensities relative to GAPDH internal loading control (n ≥ 3 independent experiments; two-way ANOVA with post hoc Tukey test; ns, not significant: p > 0.05; *p < 0.05, **p < 0.01).

Protein analysis using the total tau antibody HT7 showed positivity for distinct molecular weight bands corresponding to separate tau isoforms (Fig. 2a). Tau antibody specificity was confirmed by the recombinant human tau ladder included here as an approximate guide of the tau isoform placement as it lacks tau PTMs present in the cell lysates that could influence tau molecular weight. The comparable levels of total tau between the mutant and control iNs (Fig. S4a) also suggest that changes in pathological tau levels between the two groups are likely *MAPT* mutation-dependent and not due to changes in the overall tau levels. In agreement with previous studies showing enriched 3R tau levels in early neuronal development (41, 42), we detected predominantly 3R isoform expression in the iNs and minimal 4R tau levels, both in tau^V337M^ and tau^WT^ iNs (Fig. 2a). Antibody specificity for 3R and 4R tau was confirmed by the positivity of the respective isoform bands at the recombinant tau ladder.

Considering that oligomeric tau aggregates could represent highly toxic and pathologically significant tau species in tauopathies (43), we investigated the presence and temporal changes of tau oligomers using the antibody T18 and non-denaturing conditions for preserving the original folding state of tau (Fig. 2b). To estimate the molecular weight of the native proteins, we used a protein standard for native electrophoresis stained with Coomassie Blue for band visualization. We detected elevated oligomeric tau levels in tau^V337M^ iNs relative to controls at 2 to 3 months post differentiation that was reproducible in three independent experiments (Fig. S4 b-d). The protein standard revealed bands at 480–1,048 kD molecular weight, corresponding to tau oligomers, with no tau detected below that range. Conformational tau species identified by the MC1 antibody represent one of the most commonly detectable pathological features of tauopathies. Using the same experimental approach, we probed iN lysates using native electrophoresis. We observed a similar band pattern to T18, with elevated levels of conformational tau present after two months of culture only in the mutant iNs (Fig. 2b, S4 b-d). Overall, our results indicate a time-dependent increase of early tau toxic species in the form of oligomeric and conformationally-modified tau in the mutant iNs compared to controls at two months post differentiation, but not earlier.

Next, we examined the temporal changes in p-tau and caspase-cleaved tau levels in the iNs using the anti-phospho-tau mAb PHF1 (Ser396/404), an epitope that is phosphorylated early in tau inclusion formation in humans (44), and three caspase-cleaved tau mAbs against two neoepitope cleavage sites primarily targeted by caspase-6, D402 and D13, and the D421 site cleaved by multiple caspases, including caspases −3, −6, and −7 (8, 18). Semi-quantitative analyses of GAPDH-normalized band intensities revealed a 2.5-fold increase of PHF-1 levels in tau^V337M^ relative to controls at three months post differentiation but not earlier (Fig. 2c). Similarly, caspase-cleaved tau levels showed a 2.5 to 3-fold increase of D421 and D402-positive bands and a 2-fold increase of the D13 in tau^V337M^ iNs relative to controls at three months post differentiation. Again, these differences were absent in younger cells (Fig. 2d-f). Altogether, our results demonstrate a significantly higher accumulation of p-tau and caspase-cleaved tau in tau^V337M^ iNs relative to controls that culminated at three months post differentiation but not at earlier stages.

Next, we mapped the morphological distribution of active caspase-6 and caspase-cleaved tau markers using multiplex ICC (Fig. S5). We used three-month iNs based on our western blot analyses showing significant differences in tau pathology between mutant and controls at that time point. We detected a strong signal for active caspase-6, TauC3, mAb.D402.1, and mAb.D13.1 in the cytoplasm and neurites. Mutant and control iNs showed comparable staining intensity. Moreover, active caspase-6 positive iNs were also positive for caspase cleaved tau markers in both groups.

### Tau pathological changes in postmortem brains of FTLD and AD patients

To explore if tau pathological changes in our cell culture model are also present in the human postmortem brain, we used tissue sections from the temporal cortex of an FTLD tau^V337M^ carrier and an AD patient, both at end-stage disease. We compared these to a healthy elderly control using multiplex immunohistochemistry of active caspase-6 and caspase-cleaved tau markers (Fig. 3). In line with our *in vitro* findings, we detected active caspase-6 positivity in the cytoplasm and neurites in both cases (brown; Fig. 3). Tau^V337M^ specimens were also positive for TauC3 and mAb.D402.1 mAbs (red; Fig. 3) in the cytoplasm and neurites that co-occurred with active caspase-6. AD brain sections showed strong positivity for all caspase-cleaved tau markers, including mAb.D13.1 that was absent in FTLD, and overlapping with active caspase-6 in cytoplasm and neurites. We did not observe active caspase-6 or cleaved tau antibody positivity in brain tissue from a healthy control even after prolonged incubation with the chromogen to confirm the absence of signal (Fig. 3 g-i). Taken together, our results show that our neoepitope mAbs label pathological tau inclusions in AD and FTLD-tau and corroborate the presence of active caspase-6 in the human brain, strengthening their potential as clinically relevant therapeutic targets.

**Fig. 3.**
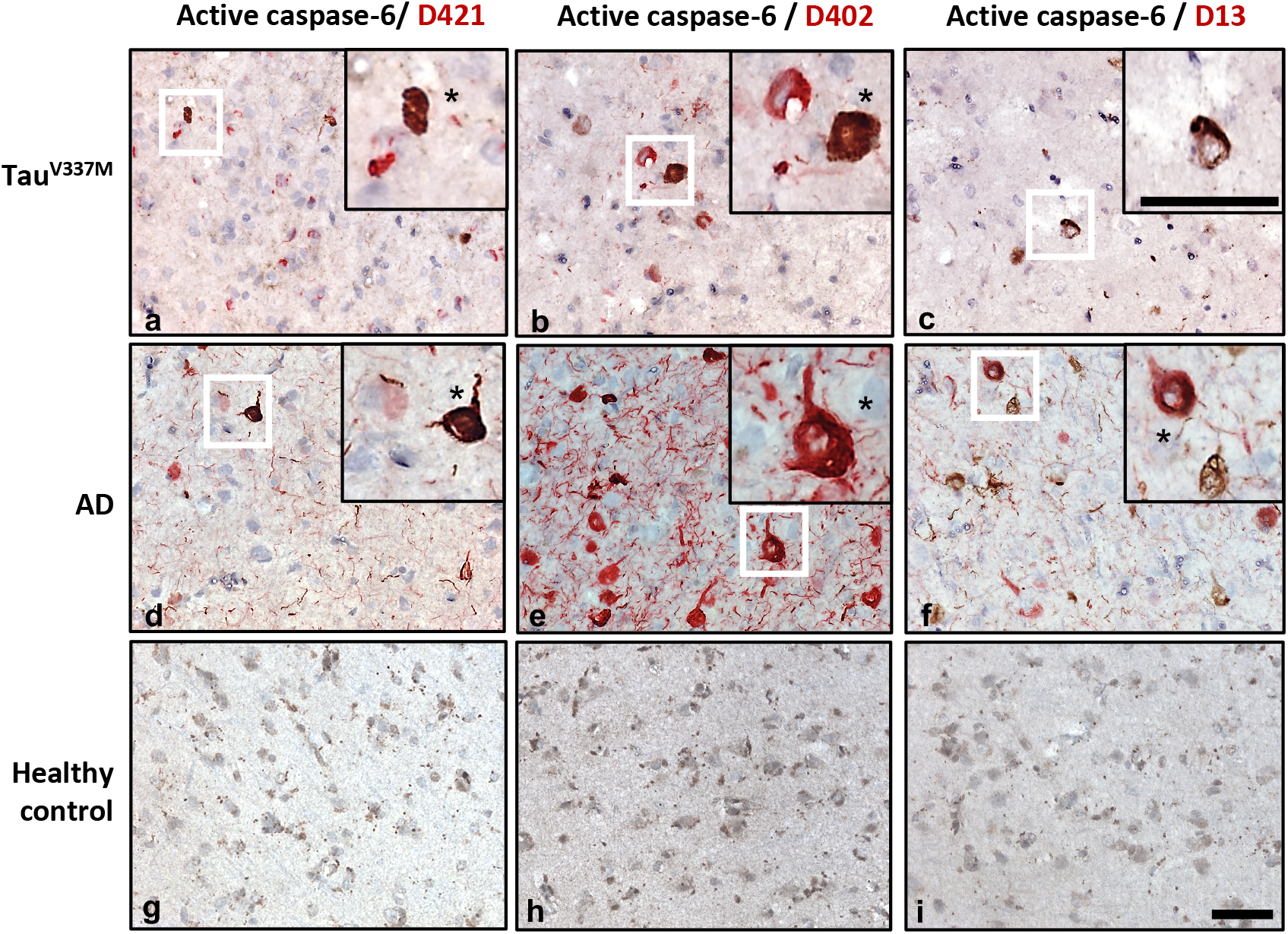
Caspase-cleaved tau antibody positivity in the postmortem brains of FTD, AD, and healthy control patients. Brain sections from the temporal cortex of an individual with tau^V337M^ and FTD (a-c), a patient with sporadic late-stage AD (d-f) showing antibody positivity for active caspase-6 (brown) and caspase-cleaved tau (red), including TauC3 (D421) (a, d), mAb.D402.1 (b, e) and mAb.D13.1 (c, f). No antibody positivity was observed in an age-match healthy control (g-i), even after prolonged incubation with the chromogen to ensure the absence of antibody signal. Insets represent the magnification of representative neurons (white squares). Asterisks indicate cells positive for both active caspase-6 and caspase-cleaved tau markers. Scale bars: 50 μm

### Increased neuronal death in the tau^V337M^ neurons

To examine if the observed accumulation of pathogenic tau in the tau^V337M^ impacts early neuronal viability, we compared the number of dead neurons between mutant and control iNs at 1-month post differentiation (Fig. 4a). We tested a panel of comparable assays commonly used for cell-death quantification, including TUNEL, Annexin-V, and ethidium homodimer III (EthD-III). However, due to high background and false-positive staining in the first two, we chose EthD-III for our quantitative analyses. Neurons positive for EthD-III, counterstained with the nuclear marker Hoechst, revealed about 60% cell death in the mutant group compared to 20% in controls, suggesting that the higher expression of pathogenic tau in tau^V337M^ iNs has a deleterious effect and leads to compromised neuronal viability (Fig. 4b).

**Fig. 4.**
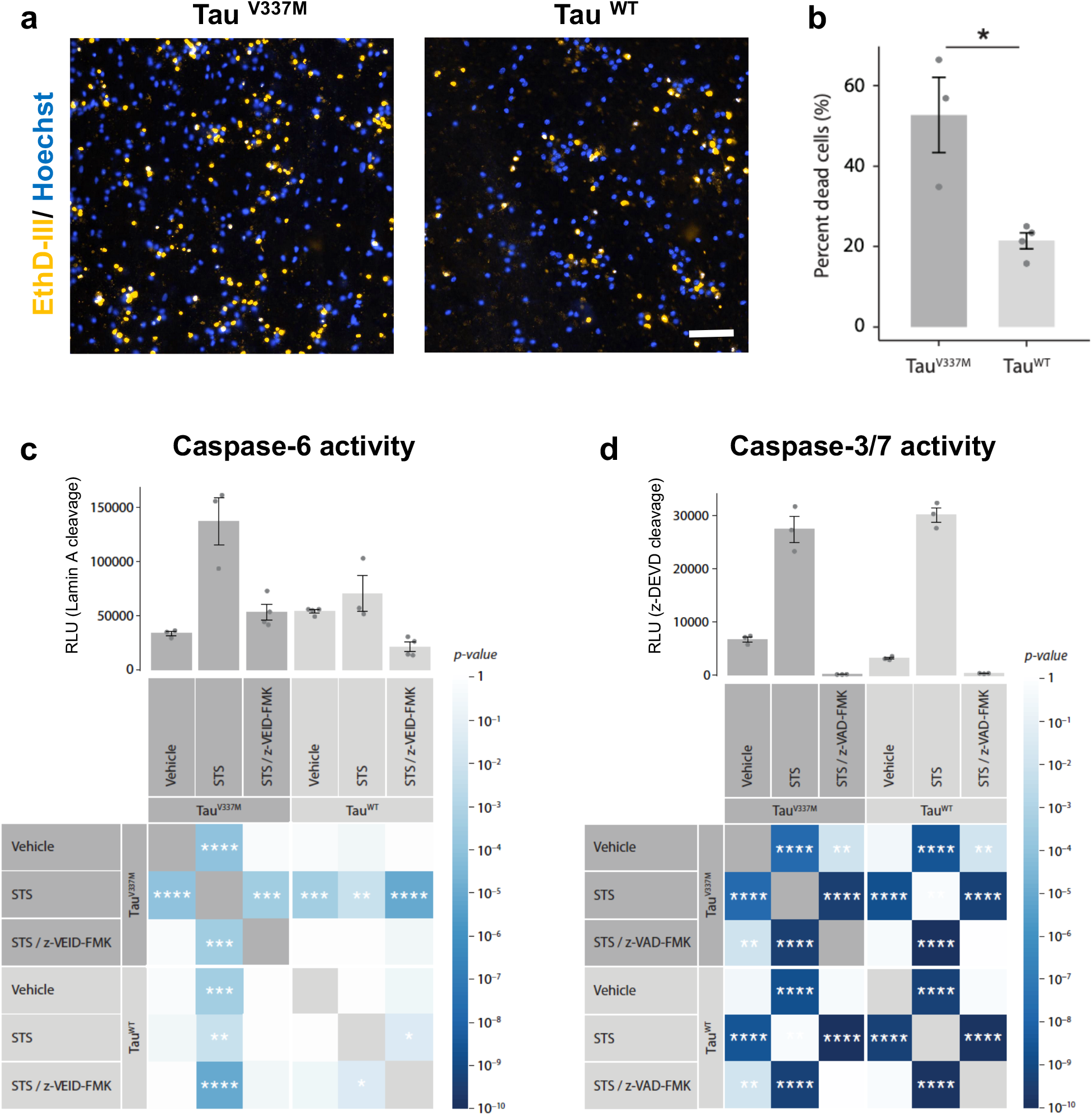
Stress-induced caspase activation and cell death in the induced neurons. (a) IF of tau^V337M^ neurons showing an increasing number of cell death detected by ethidium homodimer III staining (yellow) relative to tau^WT^ controls at 1-month post differentiation. Hoechst nuclear stain (blue) was used for the estimation of total cell numbers. (b) Quantitative analysis of neuronal death (based on a) in tau^V337M^ and control neurons. Active caspase −6 (c) and −3/7 levels (d) in 3-month neurons based on caspase-specific cleavage substrates following treatment with Vehicle (DMSO), STS, or STS and the caspase-inhibitors z-VEID-fmk (a caspase-6 inhibitor) or z-VAD-fmk (a pan-caspase inhibitor). Matrix heatmap illustrates p-values (color gradient) and significance levels (asterisks) between treatment groups. Dark gray shade represents tau^V337M^ and light gray shade represents tau^WT^ neurons (n ≥ 3 independent experiments; student t-test (b); two-way ANOVA with post hoc Tukey test (c, d); *p < 0.05, **p<0.01, ***p<0.001, ****p<0.0001). RLU: Relative light units. Scale bar: 50 μm.

### Caspase inhibition is neuroprotective against stress-induced cytotoxicity in the tau^V337M^ neurons

To test if the buildup of pathological tau in the tau^V337M^ iNs augmented their vulnerability to stressors relative to controls, we treated three month tau^V337M^ iNs with increasing concentrations of the wide-spectrum kinase inhibitor and apoptosis-inducer staurosporine (STS) for up to 48h, followed by detection of cytotoxicity levels measured by lactate dehydrogenase (LDH) release (Fig. S6). We observed a 2-fold increase of cytotoxicity levels in the tau^V337M^ iNs relative to tau^WT^ controls, using 40 μM STS for 48h (Fig. S6-b), and selected this condition for further studies. Cytotoxicity was reduced to baseline levels by treatment with the pan-caspase inhibitor z-VAD-fmk (300 μM for 48h or 4 x 75 μM in 12h intervals; Fig. S6-c), indicating that cell death occurred via apoptosis.

Based on our established conditions (Fig. S6), we exposed iNs to 40 μM STS for 48h and increasing doses of z-VAD-fmk (300 and 600 μM; Fig. 5a). Control tau^V337M^ and tau^WT^ iNs treated with vehicle (DMSO) showed comparable levels of cytotoxicity that was similar to the iNs treated only with z-VAD-fmk (600 μM) in the absence of STS (Fig. 5a). Following STS treatment, we observed almost a 5-fold increase of cytotoxicity levels in the tau^V337M^ iNs compared to a 4-fold increase in control iNs, indicating that the mutant group is significantly more vulnerable to stress (Fig. 5a). Moreover, STS co-treatment with z-VAD-fmk significantly reversed cytotoxicity levels in the mutant iNs. Next, we performed western blot analyses using lysates of iNs treated with 40 μM STS and 300 μM z-VAD-fmk. Following STS treatment, we observed a statistically significant 2-fold increase in TauC3 levels in the tau^V337M^ iNs treated with STS compared to vehicle-treated cells that was not present in control iNs (Fig. 5 b-c). In line with our cytotoxicity assay (Fig. 5a), STS co-treatment with z-VAD-fmk in the mutant iNs reversed TauC3 levels to baseline, an effect not observed in the controls.

**Fig. 5.**
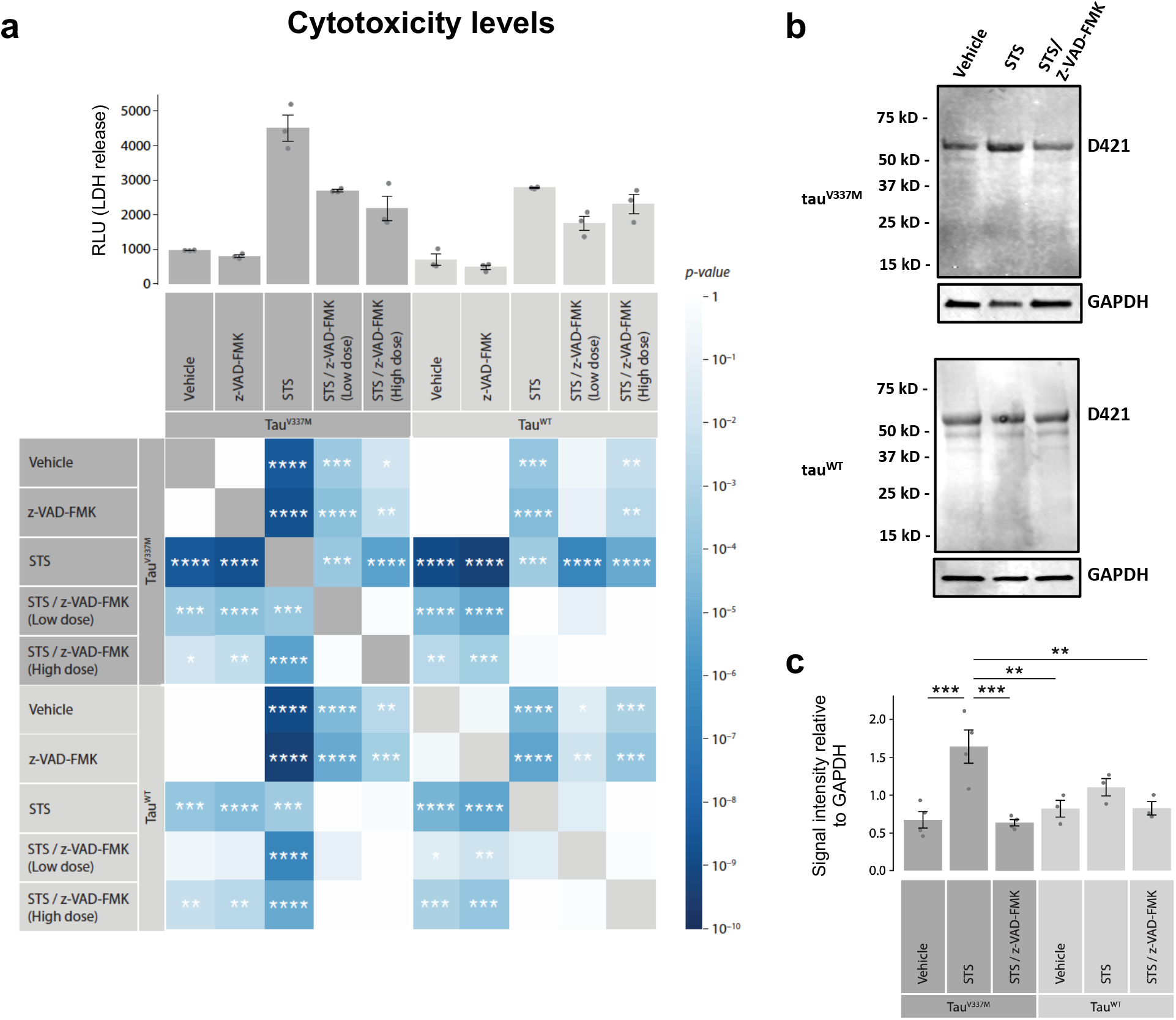
Caspase inhibition is neuroprotective against stress-induced cytotoxicity in the tau^V337M^ neurons. (A) Changes in cytotoxicity levels measured by LDH release following a 48h treatment with STS and caspaseinhibitor (z-VAD-fmk) in 12-week neurons (see Fig. S6). Matrix heatmap illustrates p-values (color gradient) and significance levels (asterisks) between treatment groups. Dark gray shade represents tau^V337M^ and light gray shade represents tau^WT^ neurons. (b-c) Semi-quantification of caspase-cleaved tau (TauC3; D421) protein levels in neurons treated with STS or STS and caspase-inhibitor for 48h based on band intensities relative to GAPDH internal loading control (n ≥ 3 independent experiments; two-way ANOVA with post hoc Tukey test; *p < 0.05, **p<0.01, ***p<0.001, ****p<0.0001). RLU: Relative light units.

Next, we assessed specific changes in active caspase −6 and −3/7 levels using a cleaved lamin A ELISA-based assay and a DEVD-aminoluciferin substrate assay, respectively (Fig. 4c-d). We observed a 4-fold increase of caspase-6 activity in the tau^V337M^ iNs compared to a 1-fold increase in control iNs following 20 μM STS treatment for 48h (Fig. 4c), indicating that the mutant group has significantly higher caspase-6 activity relative to controls. Notably, STS co-treatment with 10 μM of the caspase-6 inhibitor z-VEID-fmk significantly reversed caspase-6 activity levels in the mutant iNs. We also observed a 6-fold increase in active caspase-3/7 levels (Fig. 4d) in the tau^V337M^ and tau^WT^ treated with 40 μM STS for 6h. Caspase activity was reversed to baseline levels after the addition of 300 μM of the pan-caspase inhibitor z-VAD-fmk. Thus, STS treatment induced apoptotic cell death and a marked increase of caspase levels in the iNs that was suppressed by caspase inhibition.

Overall, our results demonstrate increased vulnerability to apoptotic cell death in tau^V337M^ iNs that was ameliorated by caspase inhibition and a strong association between tau^V337M^, caspase-mediated tau cleavage, and cytotoxicity.

### Stress-induced reduction of neurite length is rescued by caspase inhibition

The reduction in neurite length is a morphological indicator of compromised cell viability and neurotoxicity (35, 45). To examine the phenotypic effects of STS and z-VAD-fmk treatment on neurites in mutant and control iNs, we used ICC with DAPI to label cell nuclei (blue), MAP2 (red) to label cytoplasm and neurites (Fig. 6a), and automated image-based quantification of neurite length under the same treatment conditions as our cytotoxicity assay (Fig. 5a). We observed comparable mean neurite process length in untreated iNs that was significantly reduced upon treatment with 40 μM STS by 1.6 and 1.3-fold in mutant and controls, respectively (Fig. 6b). Co-treatment of STS with the z-VAD-fmk caspase inhibitor partially restored neurite length and preserved MAP2-positive processes in tau^V337M^ iNs compared to the STS-treated iNs, and fully restored neurite length in tau^WT^ iNs. These data further demonstrate that neurotoxicity following STS treatment in iNs is significantly caspase-dependent.

**Fig. 6.**
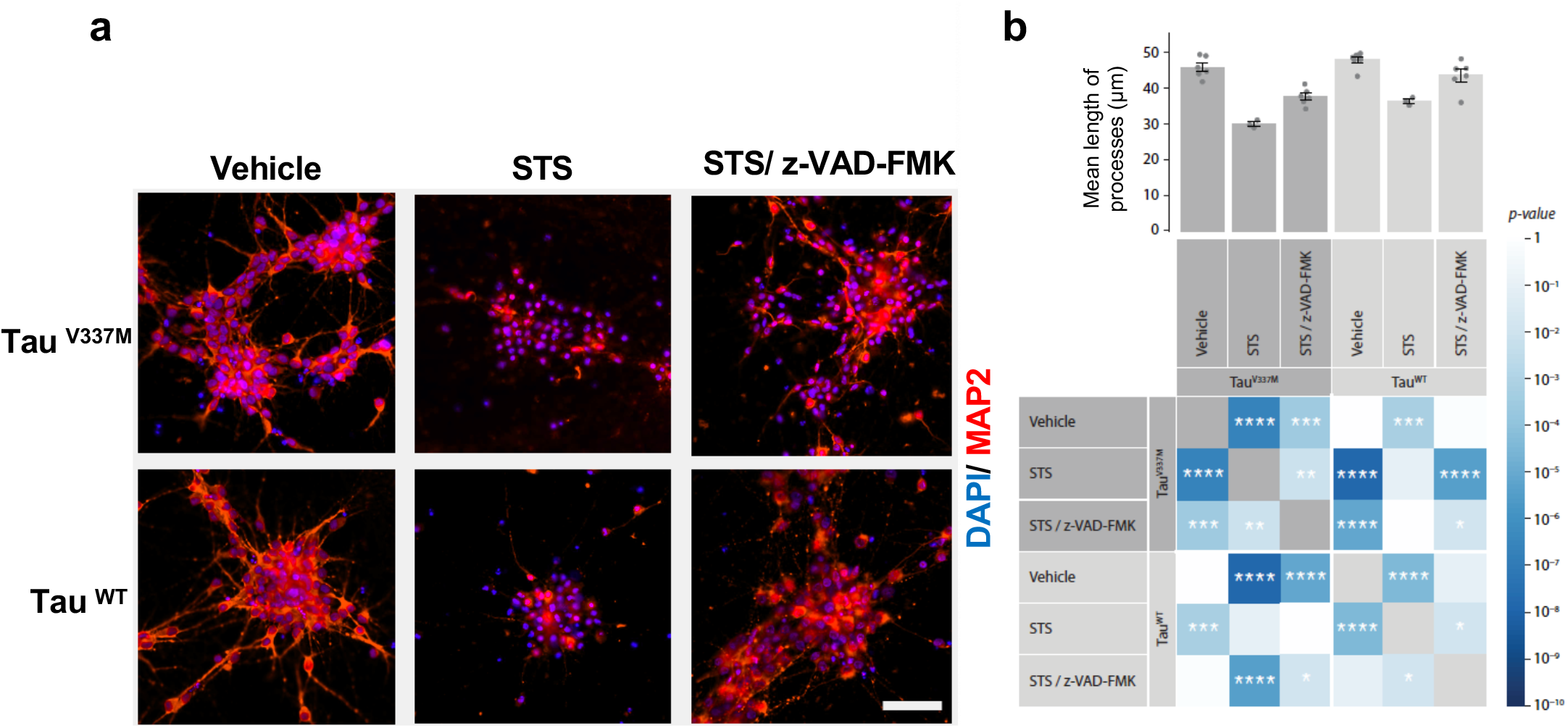
Caspase inhibition rescues stress-induced reduction of neurite length in the induced neurons. (a) IF staining of neurites (MAP2, red) and cell nuclei (DAPI, blue) following a 48h treatment with STS and caspase-inhibitor (z-VAD-fmk) in 12-week neurons. (b) Image quantification of mean length of processes (based on a). Matrix heatmap illustrates p-values (color gradient) and significance levels (asterisks) between treatment groups. Dark gray shade represents tau^V337M^ and light gray shade represents tau^m^ neurons (n ≥ 3 independent experiments; two-way ANOVA with post hoc Tukey test; *p < 0.05, **p<0.01, ***p<0.001, ****p<0.0001). Scale bar: 50 μm.

## Discussion

Active caspases can directly cleave tau, generating tau species with an enhanced propensity for self-aggregation and accumulation into neuronal cytoplasmic inclusions as proposed by biochemical studies and transgenic animal models of AD (8, 19, 28, 46). However, most of these findings are based on studies using animal models and immortalized cell lines that do not recapitulate the clinical phenotype. Building on these findings, our study innovates by using (a) novel tools for the detection of caspase-6 cleaved tau based on two mAbs against neoepitope tau cleavage sites, (b) a disease-relevant neuronal model with the FTLD-causing *MAPT* mutation tau^V337M^ that shows a progressive accumulation of pathogenic tau, including cleaved tau, p-tau, and oligomeric tau, and (c) the neuroprotective effect of caspase pharmacological intervention. Different from most studies, we maintained tau^V337M^ and tau^WT^ iNs for up to three months to avoid capturing artefactual effects caused by physiological overexpression of p-tau forms in the developing neurons (41, 47).

Three month tau^V337M^ iNs showed a three-fold increase in caspase-cleaved D402 and D13 tau protein levels and increased p-tau and oligomeric tau species compared to controls. Three month tau^V337M^ iNs also showed a significantly higher rate of neuronal death than the tau^WT^ counterpart. Treatment with staurosporine (STS), a stress inducer, led to a 5-fold increase of cytotoxicity levels in STS-treated tau^V337M^ iNs compared to vehicle-treated tau^V337M^ iNs. Although STS-induced cytotoxicity was anticipated, the increase was significantly higher in STS-treated tau^V337M^ iNs than in STS-treated control iNs. Similarly, we observed a 4-fold increase of caspase-6 activity in STS-treated tau^V337M^ iNs compared to vehicle-treated tau^V337M^ iNs that was reversed following cotreatment with the caspase-6 inhibitor z-VEID-fmk. Moreover, STS-treatment led to higher levels of caspase-cleaved D421 tau, not observed in STS-treated control iNs. Notably, co-treatment with the pan-caspase inhibitor z-VAD-fmk markedly reduced the toxic effects of STS treatment in tau^V337M^ iNs on several levels, including morphological and biochemical changes. For instance, image-based automated quantitative analyses of mean neurite length revealed a 1.6-fold length reduction of neuronal processes in tau^V337M^ iNs following STS treatment compared to 1.3-fold length reduction in controls that were markedly rescued by caspase inhibition in both lines. Altogether, our study further supports the deleterious role of caspase activation in tau pathology, the activation of caspase-6 in particular, and the therapeutic potential of caspase inhibitors against tau toxicity and cell death.

iNs with *MAPT* mutations are selectively vulnerable to stressors, likely due to increased pathogenic tau levels and a lower capacity to survive additional stress relative to WT controls (35, 38). Previous studies using tau^V337M^ iNs demonstrated progressive accumulation of toxic tau species and increased susceptibility to cell death (33, 35), a phenotype also observed in transgenic mice expressing V337M human tau that exhibit neurodegeneration and p-tau accumulation in the hippocampus, and behavioral abnormalities (48). Furthermore, tau^V337M^ iNs show tau detachment from microtubules compromised axonal transport, and dysregulation of neuronal excitability (33, 35). Our findings agree with other studies detecting tau^V337M^ iN vulnerability, including increased tau cleavage at the D421 site, immunoreactivity for the p-tau antibody AT8, and susceptibility to oxidative stress resulting in high cytotoxicity levels that were reversed by treatment with antioxidant reagents (35), strengthening the confidence of our cell lines as a model to investigate the role of caspase-cleaved tau in neurotoxicity. However, little was known about cleaved tau in this model. Here, we show an increased susceptibility of tau^V337M^ iNs to caspase-mediated tau cleavage, tau hyperphosphorylation, and stress-induced cell death relative to isogenic iNs with tau^WT^. Our novel mAbs facilitated the detection of tau species cleaved by caspase-6, an effector caspase closely associated with NFTs and neuronal loss in the human AD brain, including CA1 of the hippocampus, entorhinal cortex, and the olfactory bulb (24, 28, 30). Whereas multiple caspases target the D421 site, neoepitope sites D402 and D13 are cleaved primarily by caspase-6. These results highlight and support previous suggestions of a central role of active caspase-6 in tauopathies.

Interestingly, several of the differences detected between tau^V337M^ and tau^WT^ iNs only became apparent in neurons cultured for at least two or even three months. Western blot protein analysis of iNs cultured for 1 to 3 months revealed a time-dependent upregulation of T18 oligomeric tau and MC1 conformational tau at two months of culture in the tau^V337M^ iNs relative to controls. P-tau levels recognized by the PHF-1 antibody increased in the tau^V337M^ iNs relative to controls at three months, but not earlier. Physiological upregulation of p-tau in developing neurons is a known phenomenon *in vitro* and in human brains. Here, high levels of p-tau in tau^WT^ iNs only started to subside at two months post-differentiation. Future studies using iNs would benefit from using more mature neurons, despite the technical difficulties in maintaining neuronal lines in culture long-term. Although significantly higher levels of p-tau were only detected in three-month mutant iNs, these neurons showed higher toxicity and death as early as one month relative to controls. We can postulate that similar p-tau levels found in the two lines at one month could result in distinct cellular fates, with mutant iNs showing less tolerance relative to controls to the same levels of p-tau.

Our experimental approach offers several advantages. We introduced the heterozygous tau^V337M^ to a WT iPSC line from a healthy individual to study the pathogenic effects of the *MAPT* mutation in isolation from possible interference by the patients’ genetic background. Unlike lengthy traditional differentiation protocols that usually yield a heterogeneous population of neurons, including glia-like cells, we utilized an Ngn2-based direct differentiation protocol that improves the speed and conversion rate of iPSCs to iNs (33, 39, 40). Following this approach, we generated a homogeneous population of cortical glutamatergic neurons, a neuronal sub-type selectively vulnerable in tauopathies (49, 50), with tau^V337M^ and isogenic controls with tau^WT^ as a pre-clinical model of disease pathogenesis. Considering that early-differentiated iNs retain developmental gene expression patterns that could mask disease-specific tau changes, we cultured iNs for up to three months to ensure maturation and studied tau pathology associated with caspase activation and tau cleavage in tau^V337M^ relative to controls. Although the involvement of caspases in neurodegenerative diseases has been previously studied, our approach innovates using neoepitope mAbs against caspase-6 cleaved tau at D13 and D402 novel truncation sites. Our mAbs allowed for a rigorous examination of relevant caspase-cleaved tau species that previously could only be examined using less specific antibodies. We validated these antibodies using cellular and molecular assays and confirmed their presence in postmortem tissue from patients with AD and *MAPT* V337M FTLD-tau. Finally, the therapeutic effect of caspase inhibition against stress-induced cytotoxicity in our study supports the role of caspases as significant mediators of tau toxicity and the need for improved caspase-specific inhibitors for clinical use.

However, our approach also has limitations. Despite observing similar morphological and biochemical features in our tau^V337M^ iNs compared to other studies using tau^V337M^ lines, not all previously described changes were reproduced in our cells. For instance, we did not detect tau^V337M^-specific neurite length changes in early-differentiated iNs, as reported in another tau^V337M^ model (35). This is likely due to different neuronal induction methodologies and resulting cell phenotypes. Our approach is also limited by using peptidic, partially selective (VEID), or non-selective (VAD) caspase inhibitors currently available that prevent us from identifying conclusively which caspase(s) are responsible for tau cleavage in our cell model. Our understanding of the mechanisms of action of caspase inhibitors would tremendously benefit from follow up studies using novel, small molecule caspase −6 and −3 selective inhibitors that are currently under development. Moreover, exploration of apoptosis-related substrates targeted by caspases −3 and −6 in addition to tau, including nuclear and mitochondrial proteins, could further clarify the mechanisms of caspase-mediated neurotoxicity in our cell model.

In summary, our study demonstrates a time-dependent increase of tau cleavage and a pronounced vulnerability to stress and neuronal death in the tau^V337M^ iNs that is pharmacologically reversed by caspase inhibition. Our data support a model in which tau^V337M^ leads to activation of caspases, which in turn leads to increased vulnerability to toxic insult, including cleavage of tau to toxic or aggregation-prone tau species. While both tau^V337M^ and tau^WT^ iNs respond to STS by activating caspases, tau^V337M^ iNs are more vulnerable to tau proteolysis and cell death after caspase activation. Considering the neuroprotective effects of caspase manipulation in reducing tau cleavage and cytotoxicity and the value of differentiated human neurons as platforms for pre-clinical drug screening, future mechanistic studies on caspase-dependent pathways across *MAPT* genotypes could reveal viable therapeutic targets against tau pathogenesis in FTLD and other tauopathies.

## Materials and Methods

### Development of caspase-6 cleaved tau neoepitope monoclonal antibodies

To address the absence of mAbs against tau sites primarily cleaved by caspase-6, we generated neoepitope mAbs against cleaved tau at D402 (mAbD402; 1-402) and D13 (mAbD13; 14-441) (Fig. 1a). We produced two mAbs for each cleavage site (Table S1) by immunizing 6-8-week-old wild type Balb/c and SJL mice (SLAC) with keyhole limpet hemocyanin (KLH) conjugated tau peptides using protocols approved by ChemPartner IACUC committee. 50 μg of each peptide was injected into the abdominal cavity of each mouse along with 0.25 mL Complete Freund’s Adjuvant (Sigma). To enhance the immune response, 25 μg of KLH conjugated tau peptides was injected into the abdominal cavity of each mouse along with 0.25 mL Incomplete Freund’s Adjuvant (IFA) two weeks after the first immunization, and subsequent boosts were administered 3 weeks apart. Blood samples from each mouse were collected one week after each immunization. The antibody titer and specificity in serum were determined by enzyme-linked immunosorbent assay (ELISA) analysis against BSA-conjugated peptides I and II; and western blot against full-length tau, tau 1-402, and tau 14-441 recombinant proteins (see Supplemental Experimental Procedures).

Mice with specific immune response against tau peptides and proteins were selected for fusion and were given a final boost by intraperitoneal injection of 100 μg of the corresponding immunogen. After four days, mice were sacrificed and single-cell suspensions of splenocytes were prepared in NH4OH at 1% (w/w), followed by centrifugation at 1000 rpm and washes with DMEM (Invitrogen). Viable splenocytes were fused with mouse myeloma cells SP2/0 (ATCC) at a ratio of 5:1 with high-efficiency electric fusion (BTX ECM200). Fused cells were re-suspended in DMEM with 20% FBS and hypoxanthine-aminopterin-thymidine (HAT) medium (Invitrogen). 14 days after cell fusion, hybridoma supernatants were collected and screened by ELISA. Clones with an OD450 nm >1.0 were expanded in a 24-well plate containing DMEM with 10% heat-inactivated FBS, and supernatants were collected after 3 days culture. The antibody isotypes were determined, and ELISA and western blot were used to test their ability to bind to tau. Clones showed desired reactivity and specificity against tau were subjected to subcloning in order to get stable monoclonal hybridoma cells. Sub-cloning was carried out by limited dilution in a 96-well plate with DMEM media containing 10% FBS. Clones with specific Tau binding were further expanded in DMEM media containing 10% FBS for subsequent antibody production with Protein A purification and hybridoma cells were cryopreserved for future production. Reactivity and specificity of purified hybridoma antibodies were confirmed by ELISA against tau proteins and peptides (Fig. S1) and western blot analysis against recombinant tau proteins (Fig. S2-a, b). Considering that mAbD402.1/ mAbD402.2 and mAbD13.1/ mAbD13.2 showed comparable specificity, we used mAbD402.1 and mAbD13.1 in this study. See Supplemental Experimental Procedures for antibody specificity analysis.

### Cell lines

iPSC-derived induced neurons (iNs) with heterozygous V337M *MAPT* mutation (tau^V337M^) and WT isogenic controls (tau^WT^) were generated as previously described (33, 39). The Neurogenin 2 (Ngn2)-integrated iPSC line was created from a tau^WT^ human iPSC line (male; WTC11) (39, 51). The Tet-ON 3G-controlled Ngn2 transgene was integrated into the AAVS1 locus of human iPSC lines through a TALEN nuclease pair (39). CRISPR/Cas9 gene editing by homologous recombination was used to introduce the tau^V337M^ into the Ngn2-integrated iPSCs. Briefly, iPSCs were transfected with the Human Stem Cell Nucleofector Kit (Lonza) with sgRNA (5’-CTTGTGGGGTCA-TGGTTTACAGG-3’) plasmid (Addgene, 68463), Cas9 plasmid, and donor DNA plasmid containing a neomycin-resistance cassette (adapted from Addgene, PL552). Transfected cells were selected with neomycin for one week. Neomycin-resistant clones were selected and verified by genomic PCR and DNA sequencing. After sequence validation of the tau^V337M^ site, 1 mM Cre recombinase (Excellgen) was added to remove the neomycin-resistance cassette from gene-targeted iPSCs. Moreover, the top 10 potential off-target sites were sequenced and for each genotype the correct clones were karyotyped (Cell Line Genetics) and expanded for subsequent experiments. iNs were differentiated using a two-step protocol as previously described (33, 39). See Supplemental Experimental Procedures for detailed methods.

### Western blot protein analysis

Western blot analysis was performed using the Mini-PROTEAN Tetra system (Biorad) and standard immunoblotting techniques. See Supplemental Experimental Procedures for detailed methods.

### Cell assays and compound treatment

iNs cultured in 96-well plates were treated with either vehicle (DMSO) or staurosporine (STS) and the pancaspase inhibitor z-VAD-fmk (Enzo Lifesciences) directly in the media. Following 6h treatment, caspase-3/7 levels were examined using Caspase-Glo 3/7 (Promega) according to the manufacturer’s instructions.

An enzyme-linked immunosorbent assay (ELISA) based on lamin A cleavage, a specific substrate of active caspase-6, was used for detection and quantification of caspase-6 activity as previously described (52). Briefly, iNs were treated with either vehicle (DMSO) or staurosporine (STS) and the caspase-6 inhibitor z-VEID-fmk (Enzo Lifesciences) directly in the media for 48h. Next, cells were fixed with 4% paraformaldehyde (Thermo Fisher Scientific) in PBS for 15 min and blocked with 5% bovine serum albumin (Sigma) in PBS with 0.01% Triton X-100 for 1h at room temperature. Cells were then incubated with cleaved lamin A primary antibody (Cell Signaling, 2035, rabbit, 1:200) overnight at 4°C, followed by an HRP-conjugated secondary antibody incubation (Invitrogen) for 1h at room temperature (GE Healthcare, NA934V, donkey anti-rabbit, 1:1000) a 5 min incubation with chemiluminescent HRP substrate (Thermo Fisher Scientific), and reading with a Spectramax microplate reader (Molecular devices). iN cytotoxicity was measured using lactate dehydrogenase (LDH) release assay (Promega) following 48h treatment with vehicle (DMSO), STS, or STS/ z-VAD-fmk according to the manufacturer’s instructions. Readings were performed with a Spectramax microplate reader, followed by western blot or immunocytochemistry analysis.

To examine the morphological effects of STS and the z-VAD-fmk treatment in the iNs, we performed automated quantification of neurite length using IN Cell Developer Toolbox analysis routines based on image stacks uploaded from the IN Cell Analyzer 6500 confocal imager (GE Healthcare). To accommodate treatment-induced variation of neurite morphology during image acquisition, the Software Autofocus function was chosen to optimize the focal plane for each imaging field. DAPI was used for nuclei detection and microtubule-associated protein 2 (MAP2) for the detection of the cytoplasm and neurites. Mean neurite length per cell was calculated by subtracting a binary image of the cell body from a binary image of the entire cell, including the neurites. Data were obtained from 3-7 replicate wells per condition and an average of 1.5-2K cells per well. For IF analysis, fixed neurons were imaged using an IN Cell Analyzer 6500HS confocal imager (GE Healthcare). For further details see Supplemental Experimental Procedures.

### Human postmortem tissue processing and immunohistochemistry

Paraffin-embedded tissue sections cut at 8 μm from the temporal cortex of an individual with tau^V337M^ and FTD (68 years, female), a patient with sporadic late-stage AD and severe dementia (64 years, female, AD neuropathologic change: A3B3C3), and a healthy control free of neuropathological diagnosis (82 years, female, AD neuropathologic change: A1B1C0), were sourced from the Neurodegenerative Disease Brain Bank at the University of California, San Francisco. Immunohistochemistry was performed on de-paraffinized and rehydrated sections, following quenching of endogenous peroxidases with 3% H2O2 in methanol (Sigma) for 30 min and antigen retrieval in Tris-EDTA buffer/PBS with 0.05% Tween (Sigma) for 5 min at 121 °C in the autoclave. Sections were blocked in 5% milk/PBS with 0.05% Tween for 30 min and incubated with primary antibodies (TauC3 (D421), Invitrogen, AHB0061, mouse, 1:500; Active caspase-6 (cleaved at Asp179), Aviva Systems, OAAF05316, rabbit, 1:500; mAb.D402.1, generated in-house, clone 47G7B5, mouse, 1:500; mAb.D13.1, generated in-house, clone 5G4-1C5, mouse, 1:500) overnight at room temperature. After three washes with PBS-T (0.05% Tween-20), sections were incubated with the corresponding biotinylated secondary antibodies, followed by DAB or Red AP chromogen incubation based on manufacturer’s instructions (Vector labs) and counterstained with hematoxylin (Sigma).

### Statistics

Statistical analyses and graphics were generated using R Statistical Software (version 3.6.1; R Foundation for Statistical Computing, Vienna, Austria). Groupwise differences were analyzed by comparing means (student’s t-test, one or two-way ANOVA with post hoc Tukey tests) with the family-wise error rate set to 0.05 to correct for multiple comparisons. Data from at least three independent experiments are presented as mean ± SEM.

## Study approval

Use of human brain tissue and iPSC-derived induced neurons from humans have been approved by the University of California San Francisco Institutional Review Board.

## Author Contributions

P.T., M.R.A., and L.T.G. designed the study; P.T. performed the experiments, data acquisition and analyses. B. C.,T.Y., S. K., R.N., and M.R.A. generated the neoepitope caspase-cleaved tau antibodies. R.K. provided the oligomeric tau antibody and contributed in data interpretation. C.W. and L.G. provided the iPSC lines, C. P. contributed to data analyses and graphic design, D.B., C.W., D.O.M., C.M.K, J.E.G., and S.T. contributed with cell culture work and data interpretation, B.L.M and L.T.G provided the human brain specimens, P.T. wrote the manuscript, C.W., D.B., R.N., S.T., M.R.A., and L.T.G. revised the manuscript, L.T.G. and M.R.A. supervised the study and approved the submitted version.

## Acknowledgments

We thank Wing Hung Lee, Mifrah Hayath, Grace Pohan, Steven Chen, and Daniel Medina-Cleghorn for technical support; Dr. Andrea LeBlanc for helpful discussions on caspase biochemistry; Dr. Peter Davies for generously providing tau antibodies. This study was supported by the National Institutes of Health K01AG053433 (P.T), K24AG053435, R56AG057528 and U54 NS100717 (L.T.G), P30AG062422 and P01AG019724 (B.L.M), R01AG054025 (R.K), UCSF RAP Pilot Award program (P.T), UCSF RAP Team Science Grant (L.T.G., M.R.A.), Alzheimer’s Association AARG-16-441514 (L.T.G., M.R.A.), Rainwater Charitable Foundation (J.E.G; M.R.A.), and a Catalyst award from ShangPharma Innovation (M.R.A., T.Y., S.K, R.N.).

