## Supplemental Information for "Caspase inhibition mitigates tau cleavage and neurotoxicity in iPSC-induced neurons with the V337M *MAPT* mutation"

*** Correspondence:**

**Supplemental Experimental Procedures**

**Development of caspase-6 cleaved tau neoepitope antibodies**

**Protein expression**

Caspase-cleaved tau proteins (aa 14-441 and aa 1-402) were expressed in *E. coli*. DNA sequences of human tau were cloned into pET41a vectors with a hexa-histadine tag at the C-terminus for tau 14-441 and the N-terminus for tau 1-402. *E. coli* strain BL21(DE3) was transformed with the expression vectors and 20 mL cultures grown overnight in LB medium supplemented with 100 µg/ml kanamycin. Overnight cultures were added to 2 L LB/kanamycin medium and grown at 37°C for 3h to OD = 0.6-0.7. Tau expression was then induced with 0.2 mM IPTG at 16°C for 20 hr. Cells were pelleted, lysed, and clarified by centrifugation. Soluble protein was then loaded onto a 4 mL Ni(II) fast flow column attached to an AKTA Prime Plus FPLC. The column was washed with 100 mM Pipes, 1 mM MgSO4, pH 6.8, and tau protein eluted with 100 mM Pipes, 1 mM MgS04, 500 mM imidazole, pH 6.8. The eluant was dialyzed against a buffer of 25 mM HEPES, 0.1 mM EDTA, 0.5 mM DTT, 100 mM NaCl, pH 7.2, then loaded onto a Superdex 75 size exclusion column. Tau protein was collected at 40-43 ml. The final yield was 3.5 mg of 90% pure hexa-histadine-tagged tau.

**ELISA analysis of neoepitope antibodies**

The binding of neoepitope antibodies to antigenic tau peptides was evaluated by ELISA using 96-well microplates (Corning) coated with 100 µL of 1 µg/ml protein (full-length tau, caspase-cleaved tau, or antigen-BSA conjugate) and incubated overnight at 4°C. The plates were washed three times with PBST and blocked with 1% BSA in PBST for 2h at 37°C. Neoepitope antibodies were serially diluted in 6-fold dilutions, added to the ELISA plates, and incubated for 1h at 37°C. Plates were washed three times with PBST, and anti-mouse-HRP (1:5000; 100 µL/well; Sigma) was added for 30 min at 37°C. Plates were then washed three times with PBST, followed by 100 µL/well TMB substrate (ThermoFisher Scientific) for 30 mins, 50 µL/well 1N HCl, and absorbance measured at 450 nm using a plate reader (Molecular Devices).

**Western blot analysis of neoepitope antibodies**

Recombinant human tau was purchased from R&D (46kDa; 50 µM). Antigenic peptides were conjugated to bovine serum albumin (BSA-MBS-Tau peptide). Tau proteins and peptide-BSA conjugates were boiled in sample buffer at 90°C for 5 mins and loaded onto a 4-12% gradient Bis-Tris polyacrylamide gel in denaturing buffer (Bio-Rad). Samples and Precision Plus Protein Dual Color Standards (Bio-Rad) were electrophoresed for 2h at 100 V. Samples were then transferred to nitrocellulose membranes using iBlot2 dry transfer (ThermoFisher Scientific). Membranes were blocked in PBS/0.05% tween 20 (PBST) with 5% non-fat dry milk for 1h at room temperature and washed three times with PBST (5 mins/ wash). The primary antibody was added and incubated overnight at 4C. The membrane was then washed three times in PBST (10 mins/wash), and anti-mouse HRP (1:2000; Beyotime Biotechnology) secondary antibody added for 1h at room temperature.

The membrane was washed three times in PBST (20 mins/wash), and the signal developed by adding Beyotime ECL: mix solution A and solution B (1:1). Blots were imaged on a Bio-Rad ChemiDoc MP Imaging System with 0.5-4 sec exposures.

**Analysis of antibody specificity**

To evaluate the specificity of mAbD402.1 and mAbD13.1 to detect cleaved tau by immunofluorescence (IF), we performed antigen competition assays. mAbD402.1 and mAbD13.1 were preincubated with the peptides used as the immunogens for antibody generation (Fig. S2-c). Two immunohistochemistry experiments were run in duplicates in 96-well plates. In the primary antibody step, induced neurons (iNs) were pre-incubated with D402- or D13- immunogenic peptides, and the other two wells were untreated. All other experimental parameters remained constant. In each reaction, we also included wells in which the primary antibody was replaced by saline buffer as a negative control. We observed IF signal only in the wells incubated with the primary antibody in the absence of the cognate immunogenic peptide.

**Cell lines**

**Neuronal differentiation**

All medium, reagents, and supplements for iPSC culture and differentiation were purchased from Invitrogen (ThermoFisher Scientific), and doxycycline, dimethylsulfoxide (DMSO), cytosine β -D-arabinofuranoside (Ara-C), and poly-D-lysine from Sigma. For pre-differentiation, iPSCs were incubated with doxycycline (2 µg/mL) for 3 days at a density of 2 x $10^6$ cells/ well in six-well plates coated with Matrigel (Corning) in knockout Dulbecco's modified Eagle's medium (KO-DMEM)/ F12 medium containing N2 supplement, non-essential amino acids (NEAA), mouse laminin (0.2 mg/mL), brain-derived neurotrophic factor (BDNF, 10 ng/mL; Peprotech), neurotrophin-3 (NT3,10 ng/mL; Peprotech), and Y-27632 (Peprotech). The medium was changed daily and Y-27632 was removed from media after day 2. For maturation, precursor cells were dissociated, counted, and sub-plated in 24 or 96-well plates (50 or 25 x$10^3$ cells/ well, respectively) coated with poly-D-lysine (PDL) and laminin in maturation medium containing 50% DMEM/F12, 50% Neurobasal-A medium, 0.5 x B27 supplement, 0.5 x N2 supplement, GlutaMax, NEAA, mouse laminin (1 mg/mL), BDNF (10 ng/mL), and NT3 (10 ng/mL). Half of the medium was replaced on day 7 and again on day 14, and the medium volume was doubled on day 21. After that, one-third of the medium was replaced weekly.

**Western blot protein analysis**

iNs were harvested, washed in PBS, and lysed in N-PER buffer containing protease and phosphatase inhibitors (ThermoFisher Scientific). Lysates were sonicated using a water sonicator (Epigentek) for 5 min, followed by centrifugation at 15,000 g for 15 min at 4°C. Total protein concentration was quantified using Bradford assay (Biorad) and a Gemini XPS microplate reader (Molecular Devices). Protein lysates were mixed with 4x Laemmli buffer and incubated at 95°C for 5 minutes. Western blot analyses were performed using Mini-

PROTEAN Tetra system, 4-15% Mini-PROTEAN TGX Precast Protein Gels, dual-color precision plus protein standards (Biorad), or human recombinant tau protein ladder (Sigma). Gels were transferred onto PVDF membranes (Biorad) using standard procedures, followed by blocking in 5% nonfat milk in Tris-buffered saline with Tween 20 (TBST) for 1 h, and overnight incubation with primary tau antibodies at 4°C (PHF-1, Gift from Dr. Peter Davies, mouse, 1:1000; TauC3 (D421), Invitrogen, AHB0061, mouse, 1:1000; mAb.D402.1, generated in-house, clone 47G7B5, mouse, 1:1000; mAbD13.1, generated in-house, clone 5G4-1C5, mouse, 1:1000; T18, Gift from Dr. Rakez Kayed, rabbit, 1:1000; MC1, Gift from Dr. Peter Davies, mouse, 1:1000; HT7, Invitrogen, MN1000, mouse, 1:1000; 3R, Millipore, 05-803, mouse, 1:1000; 4R, Millipore, 05-804, mouse, 1:1000). After three washes with TBST, membranes were incubated with corresponding IRDye fluorescent secondary antibodies in intercept blocking buffer for 1h and scanned using an Odyssey Clx imaging system (Li-cor). Protein band intensities (pixel mean intensity) were quantified using Image J gel analyzer (NIH) following normalization to GAPDH (Cell signaling Technologies, 14C10, rabbit, 1:1000) endogenous control bands. For native (non-denaturing) electrophoresis, SDS and methanol were omitted, and PVDF was replaced by nitrocellulose membrane (Biorad). Unstained protein standard and Coomassie G-250 (Invitrogen) were used for molecular weight estimation of the native proteins.

**Immunocytochemistry and image analysis**

Cells were fixed with 4% paraformaldehyde (Thermo Fisher Scientific) in PBS, directly in 96-well microplates (Greiner) for 15 min and blocked with 5% bovine serum albumin (Sigma) in PBS with 0.01% Triton X-100 for 1h at room temperature. Next, cells were incubated with primary antibodies overnight at 4°C (TauC3 (D421), Invitrogen, AHB0061, mouse, 1:500; Active caspase-6 (cleaved at Asp179), Aviva Systems, OAAF05316, rabbit, 1:500; MAP2, Novus Biologicals, NB300213, chicken, 1:500; CTIP2, Abcam, ab20448, rabbit, 1:500; SATB2, Abcam, ab51502, mouse, 1:500; Vglut1, Invitrogen, 482400, Rabbit, 1:500; Nanog, Cell Signaling Technology, 4903T, Rabbit, 1:500; Oct4, Cell Signaling Technology, 2840T, Rabbit, 1:500; Sox2, Cell Signaling Technology, 3579T, Rabbit, 1:500; mAb.D402.1, generated in-house, clone 47G7B5, mouse, 1:500; mAb.D13.1, generated in-house, clone 5G4-1C5, mouse, 1:500). After three washes with PBS-T (0.05% Tween-20), cells were incubated with the corresponding secondary antibodies conjugated with Alexa Fluor 488, 555 or 647 (Invitrogen) for 1h at room temperature, followed by three washes with PBS-T, and 4′,6-Diamidino-2-phenylindole dihydrochloride (DAPI; Sigma) counterstaining for nuclei identification. Cells were stored submerged in PBS at 4°C until imaging. For cell-death detection, unfixed cells were incubated with Ethidium Homodimer III, followed by Hoechst 33342 at room temperature for 15 minutes in the dark according to the manufacturer’s instructions (Biotium). Multiplex immunofluorescent (IF) images were acquired directly from the 96-well plates using an IN Cell Analyzer 6500HS confocal imager (GE Healthcare) with a 20x/0.75 objective (Nikon Plan Apo).

### Supplemental Figures and Legends

**Fig. S1** Determining selectivity of tau neoepitope monoclonal antibodies using ELISA. (a) Mouse monoclonal antibodies (mAbs) mAbD402.1 (red circles) and mAbD402.2 (orange squares) binding to tau (1-402). (b) mAbD402.1 (red circles) and mAbD402.2 (orange squares) binding to full-length tau (1-441). (c) mAbD13.1 (red circles) and mAbD13.2 (orange squares) binding to tau (14-441). (d) mAbD13.1 (red circles) and mAbD13.2 (orange squares) binding to full-length tau (1-441). (e) mAbD13.1 (red circles) and mAbD13.2 (orange squares) binding to BSA conjugated to peptide I ($H^{14}AGTYGLGDRK^{24}C$). (f) mAbD13.1 (red circles) and mAbD13.2 (orange squares) binding to BSA conjugated to peptide II ($CI^{392}VYKSPVVSGD^{402}$). See also Table S1. NC (green triangles): no protein immobilized.

**Fig. S2** Determining selectivity of tau neoepitope monoclonal antibodies using western blot protein analysis and IF. (a) Mouse monoclonal antibodies (mAbs), mAbD402.1 and mAbD402.2 bind to tau truncated at D402 (lanes 1, 3, 6, 7) but not full-length tau (lanes 2, 4, 7, 9). (b) mAbD13 binds to tau 14-441 (lanes 4, 9) and BSA conjugated to peptide I ($H^{14}AGTYGLGDRK^{24}C$; lanes 2, 7) but not full-length tau (lanes 1, 6) or BSA conjugated to peptide II ($CI^{392}VYKSPVVSGD^{402}$; lanes 3, 8). Lane 5 (a, b) contains the molecular weight markers. (c) IF multiplex staining of cell nuclei using DAPI (blue), caspase-cleaved tau at D402 and D13 epitopes (green), active caspase-6 (orange), and neuronal marker MAP2 (red) in 4-weeks $tau^{V337M}$ iNs. Cells without immunogen peptide treatment (see Table S1) are positive for mAbD402.1 and mAbD13.1 (green; upper panel). Incubation with the immunogen peptides diminished antibody positivity (green; bottom panel). Scale bar: 60 μm.

**Fig. S3** Characterization of the $tau^{V337M}$ and $tau^{WT}$ iPSC lines. (a) G-banded karyotype analysis of $tau^{V337M}$ and $tau^{WT}$ iPSCs. (b) Sequencing from the genomic DNA of the iPSC lines confirmed the presence of the heterozygous $tau^{V337M}$ in exon 12. (c) IF staining for the pluripotency markers NANOG, OCT4, and SOX2 (green). Nuclei were labeled with DAPI (blue). Scale bar:100 μm.

**Fig. S4** Changes in total, oligomeric, and conformational tau levels in neurons cultured for 1 to 3 months. Semi-quantification of total tau protein levels (a) detected by the antibody HT7 based on band intensities relative to GAPDH internal loading control (see also Fig. 4a). Western blots from three independent experiments (b-d) were performed under non-denaturing conditions and probed for oligomeric (T18; top panels) and conformational tau (MC1; bottom panels). A protein standard for native electrophoresis stained with Coomassie blue (far-left panels) was used for estimating the molecular weight of the proteins ($n \geq 3$ independent experiments; two-way ANOVA with post hoc Tukey test; ns, not significant).

**Fig. S5** Tau pathology in the induced neurons. Multiplex IF of caspase-cleaved tau (D421, D402, and D13 epitopes; green), caspase-6 (orange), and the neuronal marker MAP2 (red) in $tau^{V337M}$ (a) and $tau^{WT}$ (b) neurons cultured for three months. Nuclei were counterstained with DAPI (blue). Scale: 60 µm.

**Fig. S6** Establishing treatment conditions for cytotoxicity analyses in the induced neurons. Differences in cytotoxicity levels between $tau^{V337M}$ and $tau^{WT}$ neurons were based on LDH release after 24h (a) and 48h (b) treatment with increasing STS doses. (c) Changes in cytotoxicity levels in $tau^{V337M}$ iNs after treatment with 40 µM STS and the caspase inhibitor z-VAD-fmk as a single dose (300 µM) or in sub-doses (4 x 75 µM) at 12h intervals. Matrix heatmap illustrates p-values (color gradient) and significance levels (asterisks) between treatment groups. Dark gray shade represents $tau^{V337M}$ neurons (n = 3 independent experiments; *p < 0.05, **p<0.01, ***p<0.001, ****p<0.0001). RLU: Relative light units.

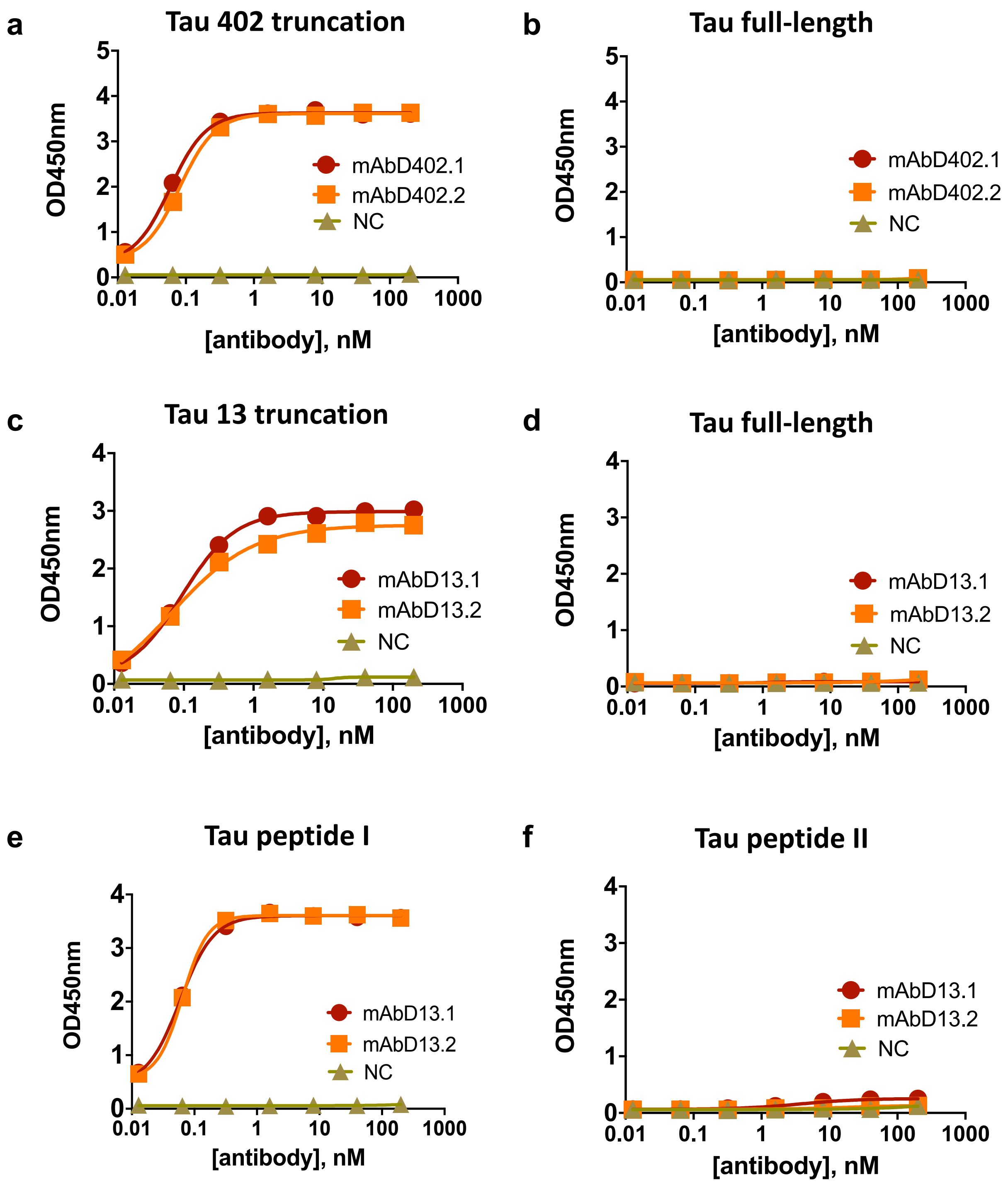

a
Tau 402 truncation
OD450nm
[antibody], nM
mAbD402.1
mAbD402.2
NC
b
Tau full-length
OD450nm
[antibody], nM
mAbD402.1
mAbD402.2
NC
c
Tau 13 truncation
OD450nm
[antibody], nM
mAbD13.1
mAbD13.2
NC
d
Tau full-length
OD450nm
[antibody], nM
mAbD13.1
mAbD13.2
NC
e
Tau peptide I
OD450nm
[antibody], nM
mAbD13.1
mAbD13.2
NC
f
Tau peptide II
OD450nm
[antibody], nM
mAbD13.1
mAbD13.2
NC


**Fig. S1**

**a**

| Lane | 1 | 2 | 3 | 4 | 5 | 6 | 7 | 8 | 9 |
|---|---|---|---|---|---|---|---|---|---|
| tau | 1-402 | 1-441 | 1-402 | 1-441 | MW | 1-402 | 1-441 | 1-402 | 1-441 |
| 1° | mAbD 402.1 | mAbD 402.1 | mAbD 402.2 | mAbD 402.2 | | mAbD4 02.1 | mAbD 402.1 | mAbD 402.2 | mAbD 402.2 |
| concen tration | 10 μg/mL | | | | | 1 μg/mL | | | |

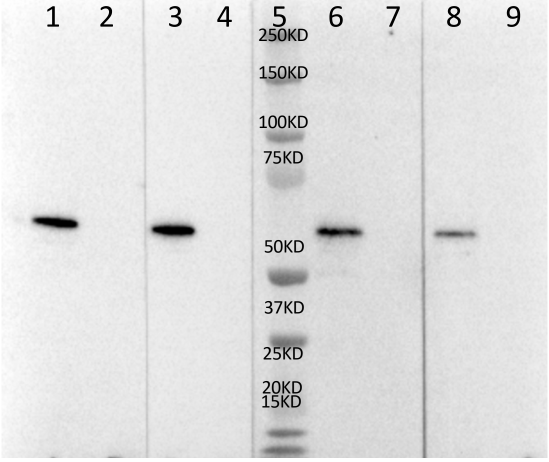


**b**

| Lane | 1 | 2 | 3 | 4 | 5 | 6 | 7 | 8 | 9 |
|---|---|---|---|---|---|---|---|---|---|
| tau | 1-441 | BSA-pepI | BSA-pepII | 14-441 | MW | 1-441 | BSA-pepI | BSA-pepII | 14-441 |
| mAbD 13.1 | 10 μg/mL | | | | | 1 μg/mL | | | |

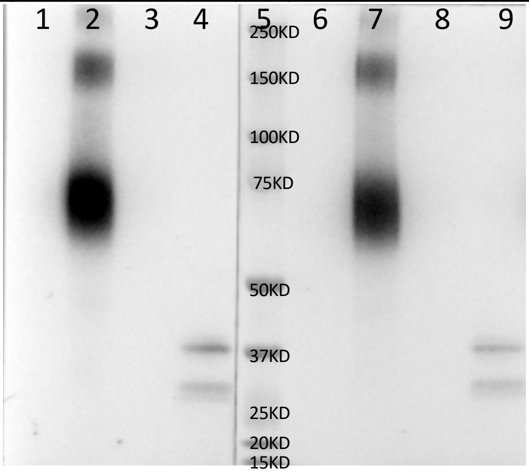


**c**

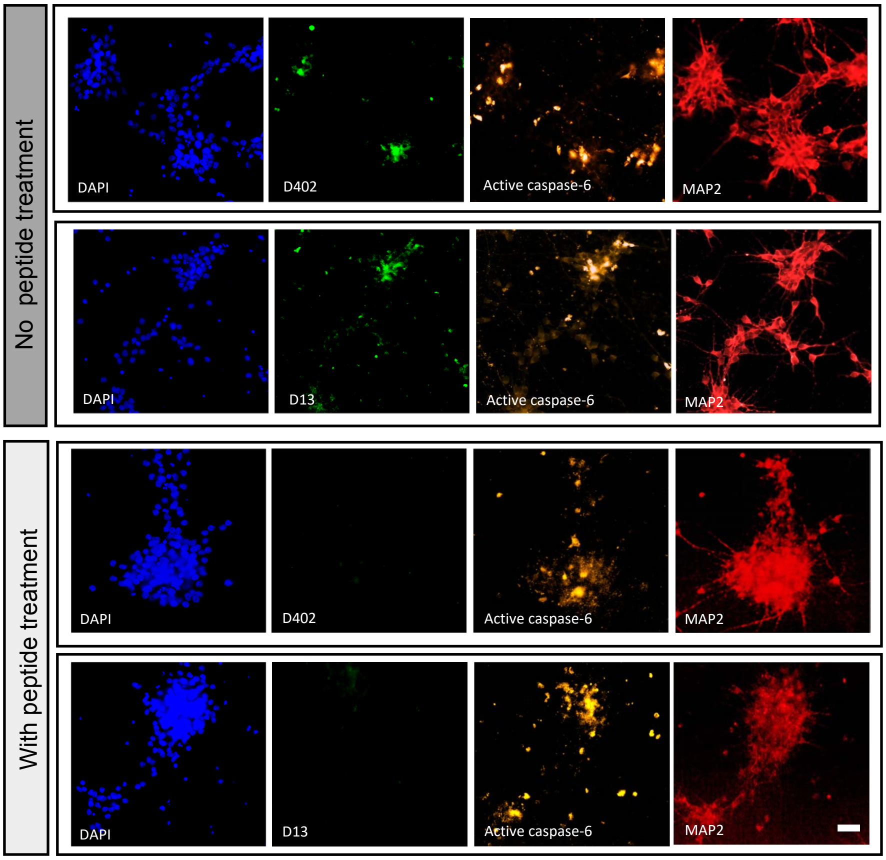


**Fig. S2**

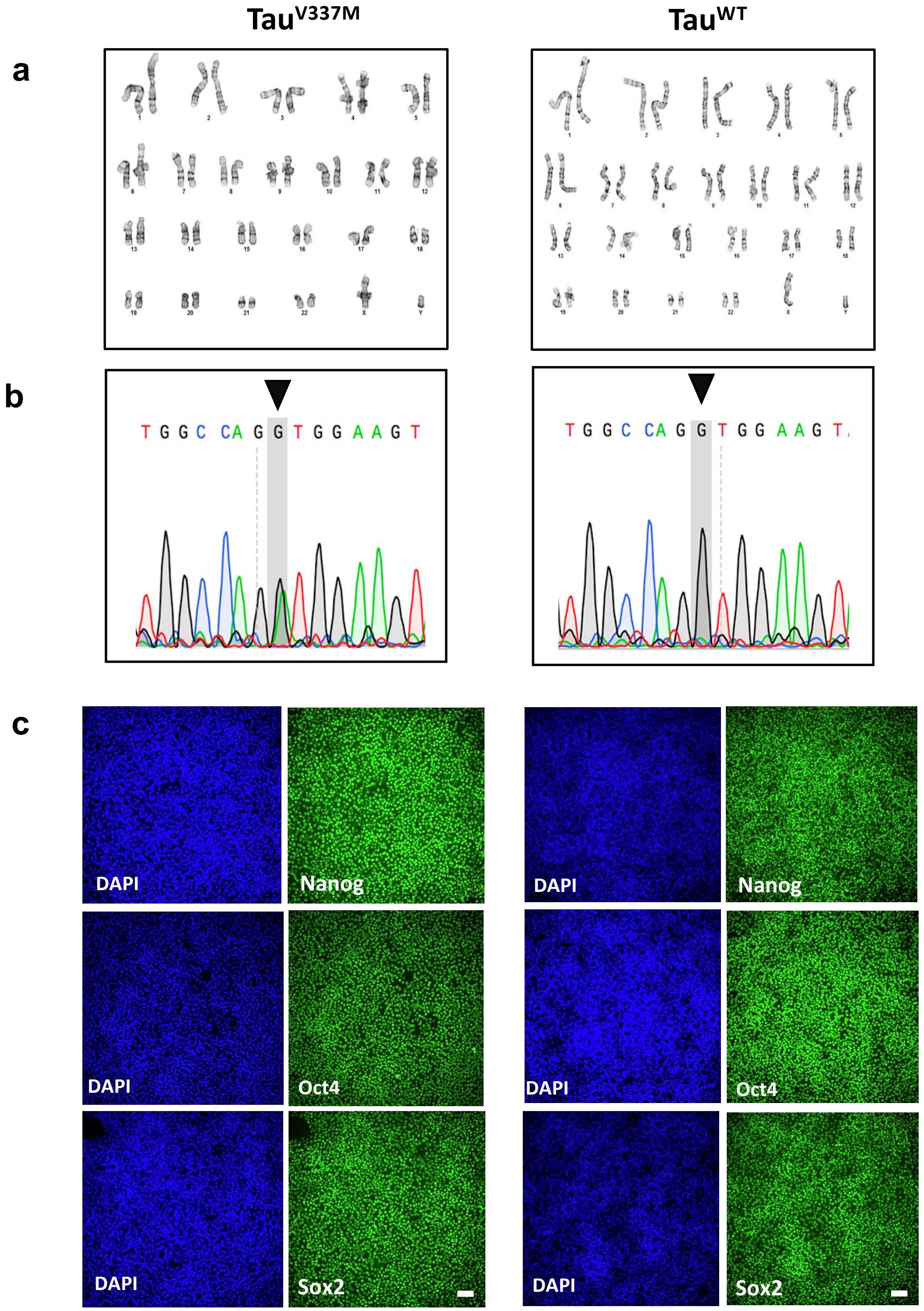
Tau$^{V337M}$
Tau$^{WT}$
a
b
T G G C CA G G T G G A A G T
T G G C CA G G T G G A A G T
c
DAPI
Nanog
DAPI
Oct4
DAPI
Sox2
DAPI
Nanog
DAPI
Oct4
DAPI
Sox2

Fig. S3

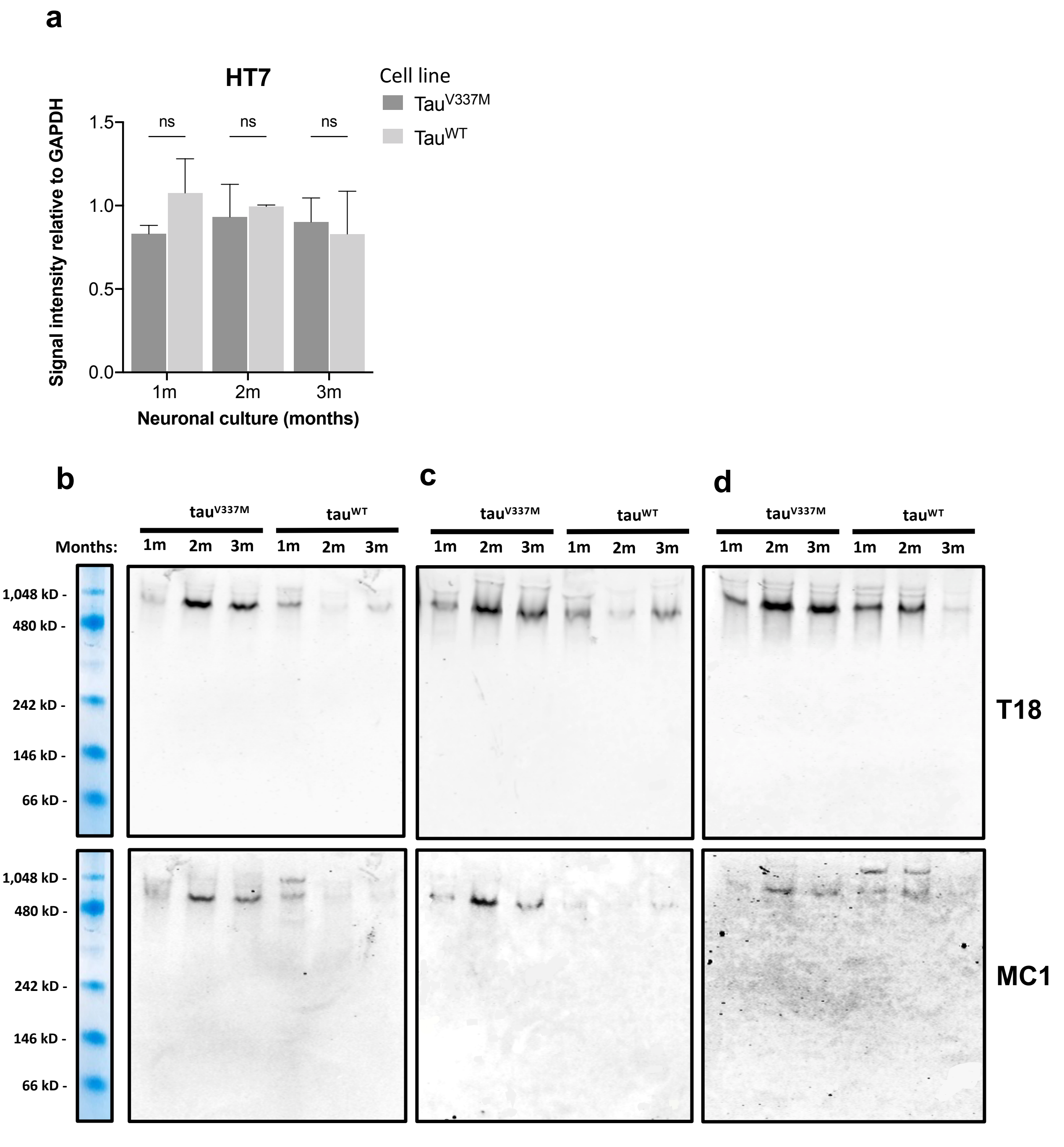

a
HT7
Cell line
TauV337M
TauWT
Signal intensity relative to GAPDH
1.5
1.0
0.5
0.0
ns
ns
ns
1m
2m
3m
Neuronal culture (months)
b
c
d
tauV337M
tauWT
Months:
1m 2m 3m 1m 2m 3m
1,048 kD -
480 kD -
242 kD -
146 kD -
66 kD -
T18
MC1


Fig. S4

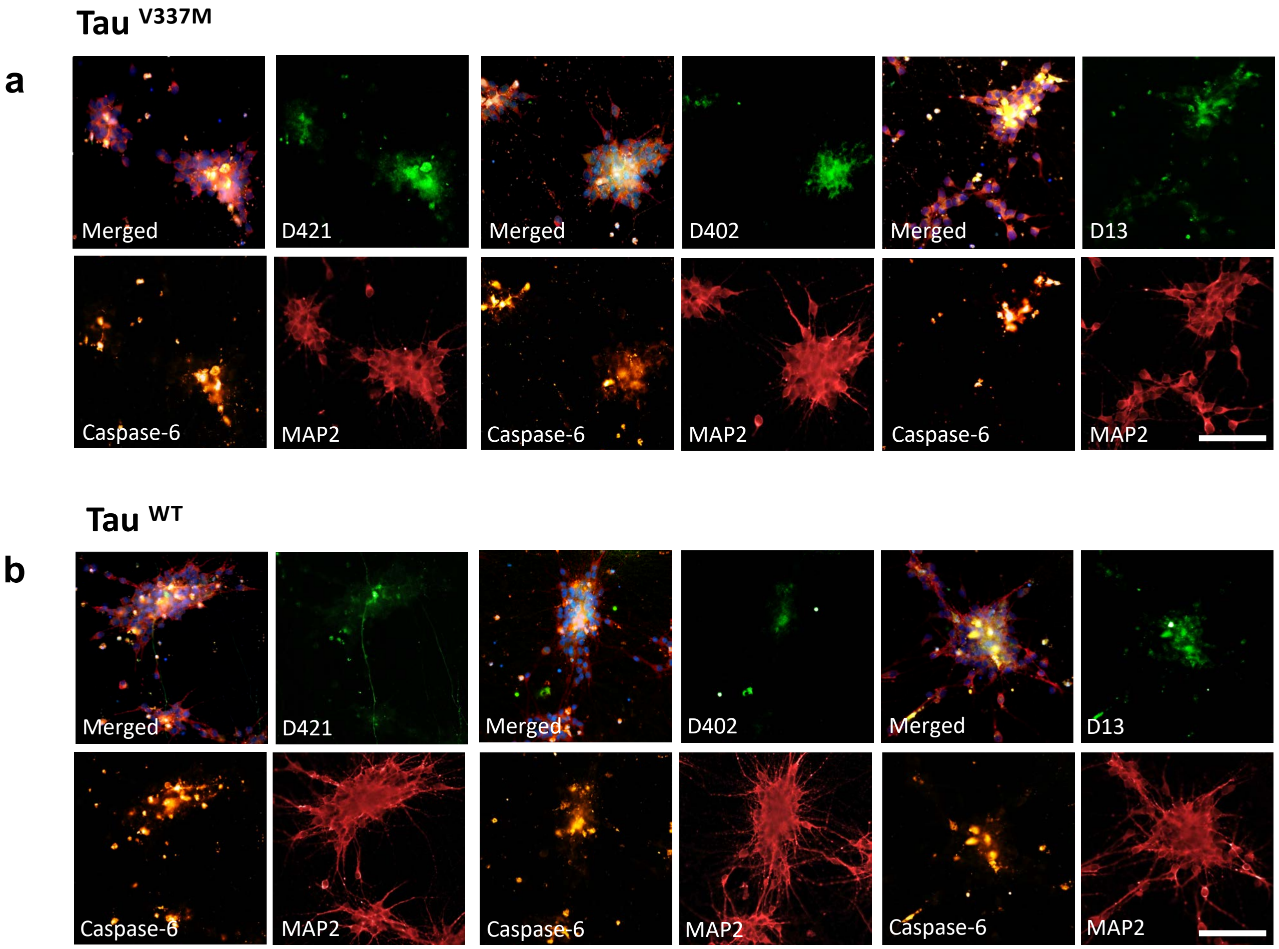
Tau V337M
a
Merged
D421
Merged
D402
Merged
D13
Caspase-6
MAP2
Caspase-6
MAP2
Caspase-6
MAP2
Tau WT
b
Merged
D421
Merged
D402
Merged
D13
Caspase-6
MAP2
Caspase-6
MAP2
Caspase-6
MAP2

**Fig. S5**

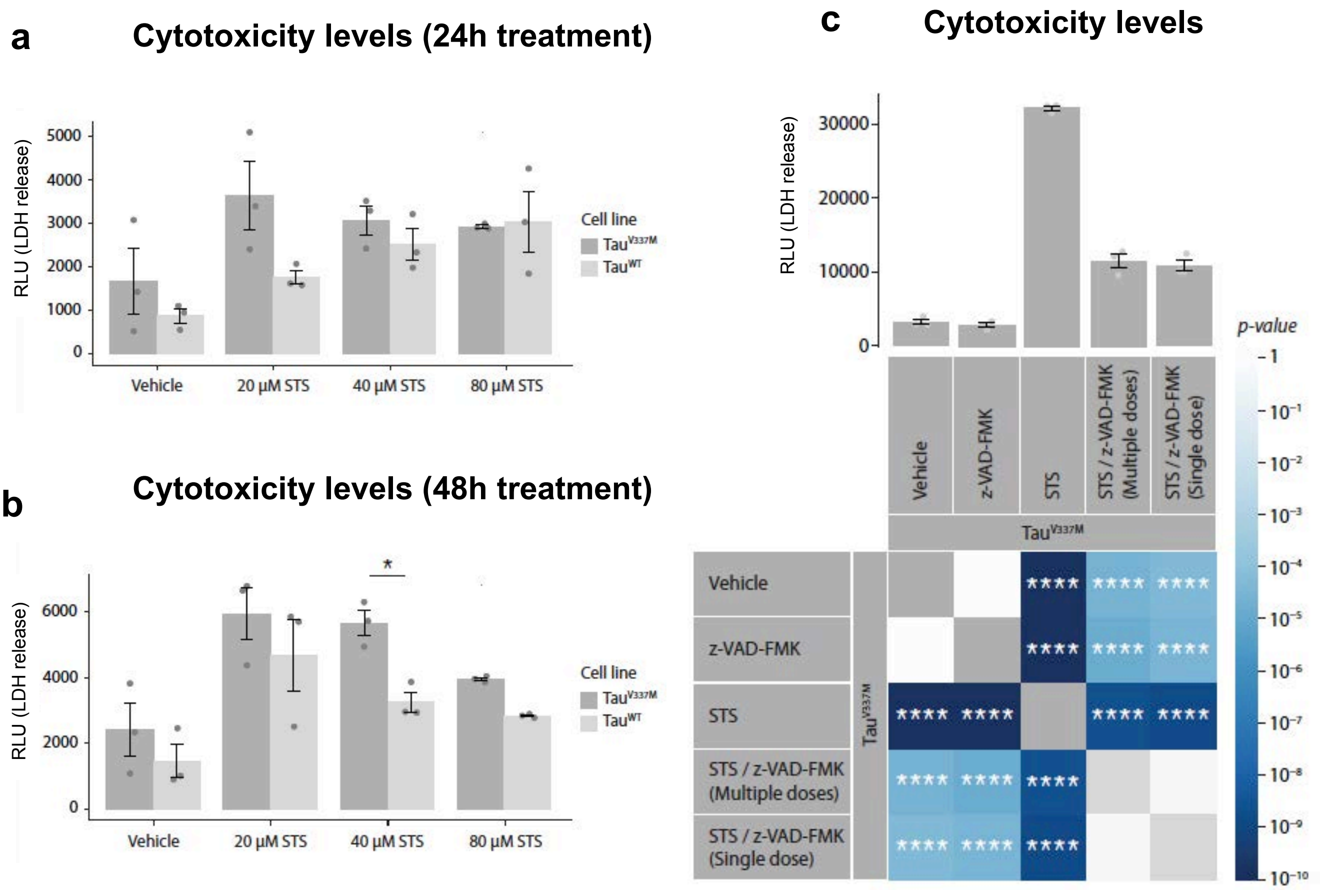
a
Cytotoxicity levels (24h treatment)
RLU (LDH release)
5000
4000
3000
2000
1000
0
Vehicle
20 µM STS
40 µM STS
80 µM STS
Cell line
Tau$^{V337M}$
Tau$^{WT}$
b
Cytotoxicity levels (48h treatment)
RLU (LDH release)
6000
4000
2000
0
*
Vehicle
20 µM STS
40 µM STS
80 µM STS
Cell line
Tau$^{V337M}$
Tau$^{WT}$
c
Cytotoxicity levels
RLU (LDH release)
30000
20000
10000
0
Vehicle
z-VAD-FMK
STS
STS / z-VAD-FMK (Multiple doses)
STS / z-VAD-FMK (Single dose)
Tau$^{V337M}$
Vehicle
z-VAD-FMK
STS
STS / z-VAD-FMK (Multiple doses)
STS / z-VAD-FMK (Single dose)
Tau$^{V337M}$
****
p-value
1
$10^{-1}$
$10^{-2}$
$10^{-3}$
$10^{-4}$
$10^{-5}$
$10^{-6}$
$10^{-7}$
$10^{-8}$
$10^{-9}$
$10^{-10}$

**Fig. S6**

**Table S1. Anti-tau neoepitope antibodies**

| Antibody | Immunogen | Conc (mg/ml) | Volume (ml) | Quantity (mg) |
|---|---|---|---|---|
| mAbD402.1 | KLH-CIVYKSPVVSGD | 0.861 | 11 | 9.47 |
| mAbD402.2 | KLH-CIVYKSPVVSGD | 0.307 | 14 | 4.3 |
| mAbD13.1 | HAGTYGLGDRKC-KLH | 1.109 | 13 | 14.417 |
| mAbD13.2 | HAGTYGLGDRKC-KLH | 1.387 | 10 | 13.87 |

mAb: monoclonal antibody
